# S-nitrosylation of protein kinase A is required for its activation by GPCRs

**DOI:** 10.64898/2026.05.27.727026

**Authors:** Sherif Bahriz, Zhenduo Zhu, Bing Xu, Jian Wu, Ning Cao, Chaoqun Zhu, Wen Pan, Caeser Tawfeeg, Zhuer Zeng, Johannes W. Hell, Qingtong Wang, Richard Premont, Ying Wang, Jonathan S. Stamler, Susan S. Taylor, Yang K. Xiang

## Abstract

Stimulation of many G protein-coupled receptors (GPCRs) increases cyclic adenosine monophosphate (cAMP) and nitric oxide (NO). While cAMP-dependent activation of protein kinase A (PKA) is a central regulatory mechanism, a parallel role for NO in GPCR transduction has not been established. Here we show that upon stimulation of multiple GPCRs in heart, brain, and fat, the regulatory subunits of PKA undergo enzymatic S-nitrosylation by SNO-CoA-assisted nitrosylase (SCAN). S-nitrosylation by SCAN is required for dissociation of PKA holoenzymes and activation of PKA. In transgenic and cardiovascular disease models, impaired adrenergic stimulation is identified with deficient S-nitrosylation of PKA, which can be rescued by the drug sodium nitroprusside. Our work suggests a new understanding of GPCR physiology with direct application to the clinical setting.

## Main

GPCRs are the most important cell surface sensors responding to hormones and neurotransmitters, and represent the predominant family of drug targets (1). The largest class of GPCRs couples to the stimulatory heterotrimeric G protein (Gs) to regulate a broad range of physiological responses in cognitive behaviors, cardiovascular function, and metabolism (2,3). In particular, ligand stimulation of the GPCR-Gs cascade leads to activation of adenylyl cyclases that increase production of the intracellular messenger cyclic adenosine monophosphate (cAMP). One major effect of elevated cAMP is binding regulatory subunits of protein kinase A (PKA) holoenzymes to release the active PKA kinase catalytic subunit to phosphorylate target proteins mediating physiological responses ranging from neuronal excitation to muscle contraction, hormone secretion and fuel metabolism across numerous tissues (4–7). Conversely, the GPCR-induced cAMP-PKA pathway is commonly impaired in disease, motivating therapeutic approaches in conditions ranging from depression and heart failure to obesity (8,9).

Stimulation of GPCRs is also known to activate nitric oxide synthases (NOSs), which produce the secondary messenger, nitric oxide (NO) (10,11). Mechanistically, NO induces physiological responses via S-nitrosylation, the direct post-translational modification of cysteine residues in target proteins (12,13), as well as by activating soluble guanylyl cyclase (sGC) to increase production of the intracellular messenger cyclic guanosine monophosphate (cGMP) that activates protein kinase G (PKG) to phosphorylate downstream effectors (14–16). The canonical role of the NO-cGMP-PKG cascade is in vessel relaxation (17,18), which is often impaired in cardiovascular diseases (19,20) and has been successfully targeted in heart failure, erectile dysfunction, and pulmonary hypertension (21,22). On the other hand, S-nitrosylation broadly affects proteins of all classes (over 10,000 proteins are reported to be S-nitrosylated), including many GPCRs, most signaling machinery and downstream effectors (23,24). Thus, an emerging picture suggests regulation of GPCR signaling by both S-nitrosylation and phosphorylation and raises the idea of integrated effects.

The prototypic β₂-adrenergic receptor (β₂AR) provides an illustrative case. Receptor stimulation promotes site-specific S-nitrosylation at Cys265 in parallel with PKA-mediated phosphorylation at Ser261/262. These co-incident modifications are both necessary for the classic desensitization of this GPCR by PKA, revealing an essential role for S-nitrosylation in canonical actions of PKA (25). More recently, the enzyme S-nitroso-coenzyme A-assisted nitrosylase (SCAN), encoded by the biliverdin reductase B (BLVRB) gene, has been shown to carry out targeted S-nitrosylation in receptor systems, including receptor tyrosine kinases (RTKs) such as insulin receptor, highlighting enzymatic convergence between RTK and GPCR pathways that mirrors kinase-mediated crosstalk through GPCR receptor kinases (GRKs) or the proto-oncogene tyrosine-protein kinase Src (26,27). In these studies, S-nitrosylation presents as an emerging paradigm as a reversible, site-specific regulatory switch that integrates NO production with phosphorylation-dependent signaling fidelity in both receptor classes.

Notwithstanding the above, NO has been largely ignored in canonical GPCR signaling. In this study, we identify an essential role for S-nitrosylation in activation of the paradigmatic GPCR effector PKA, in brain, heart and fat tissues. Sites of S-nitrosylation in PKA are conserved within the C-terminal cAMP-binding domain (CNB-B) of PKA regulatory subunits across families and species. We further show that SCAN is necessary for S-nitrosylation and consequent activation of PKA stimulated by a wide range of different GPCRs, including adrenergic, prostaglandin, glucagon-like peptide 1 (GLP1), glucose-dependent insulinotropic polypeptide (GIP), adenosine, and dopamine receptors. In brain, heart and fat tissues lacking either NOS1 or NOS3, or expressing PKA mutants refractory to S-nitrosylation, GPCR stimulation of PKA activity is impaired. Further, in a heart failure model, where β1 adrenergic receptor (β1AR)-NOS1-PKA coupling was found to be reduced, the clinically-used vasodilatory NO donor drug sodium nitroprusside (SNP) effectively rescued PKA activity, substrate phosphorylation, excitation-contraction coupling, and cardiac contractility.

Taken together, our study defines a new paradigm for GPCR signaling where NO is central to canonical PKA as well as PKG physiology, thus revising understanding of a wide range of neuronal, cardiovascular, and metabolic disorders.

## RESULT

### GPCR activation of PKA requires NO production

The role of NO in the β-adrenergic receptor-cAMP-PKA signaling pathway was first examined through cardiac phosphoproteome profiling in heart lysate after stimulation of wildtype and NOS1-KO mice with the clinically used β1AR-selective agonist dobutamine (DOB). The heat map shows a marked reduction in phosphorylation of PKA target proteins in NOS1-KO hearts compared with WT controls, and annotation confirmed a global loss of PKA substrate phosphorylation in muscle contraction and cytoskeleton proteins across the NOS1-KO samples (Figure 1A; fig S1A-1D). Phosphoproteomic change was confirmed by Western blotting for phospholamban, a canonical PKA substrate involved in cardiac excitation-contraction coupling. DOB-induced phosphorylation of phospholamban at Ser16 (pPLB16) was attenuated in isolated NOS1-KO in adult ventricular myocytes (AVMs, Figure 1B). Having established the critical role of NOS1 in activation of PKA in heart, we extended our interrogation to brain and brown adipose tissue (BAT) to test whether NOS-dependent activation of PKA activity is a shared mechanism across various tissues. Using ELISA-based assays, we measured cAMP, cGMP, and PKA activity in tissue lysates following receptor-specific stimulation with DOB and SKF81297 (SKF, a dopamine D1 receptor agonist) for heart and brain, and CL316,243 (CL, a β3AR-adrenergic receptor selective agonist) for BAT. Agonist stimulation increased cAMP, cGMP, and PKA activity across all three WT tissues. In NOS1-KO heart and brain, cAMP levels rose to the same extent as in WT, but cGMP level, an indirect readout of NO availability, was blunted as expected in the absence of the major NOS isoform. Notably, PKA activity was coincidentally diminished (Figure 1C, D; fig S1E-1G). While CL treatment elevated cAMP, cGMP, and PKA in WT BAT, NOS3-KO BAT showed preserved cAMP but impaired cGMP and PKA activation, in agreement with the known β3AR-NOS3 coupling in adipose tissue (Figure 1E). Consistent with blunted PKA activity, the PKA phosphorylation of substrates induced by these GPCR ligands were reduced in NOS deficient tissues (fig S2 A-D) These results demonstrate that NOS isoform-medidated production of NO is essential for coupling GPCR-driven cAMP signaling to full PKA activation and phosphorylation across multiple tissues, and that PKA activation is not accounted for solely by cAMP level. To extend these tissue-level observations to whole-organ physiology, we assessed cardiac function by echocardiography. In WT mice, DOB increased cardiac ejection fraction (EF), whereas this response was markedly blunted in NOS1-KO but preserved in NOS3-KO mice, indicating a selective requirement for NOS1 in β1AR inotropy (Figure 1F; fig S3A-D; table S1-S2). However, the non-selective β1AR/β2AR agonist isoproterenol (ISO) augmented EF across genotypes, including NOS1-KO, and β1AR-KO, consistent with compensatory β2-adrenergic signaling (fig S3E-G). Moreover, we subjected transgenic mice lacking either protein kinase G1 (PKG1), PKG2, or both to challenge with DOB. Deleting PKG isoforms did not affect DOB induced increases in EF (fig S4A, B; table S3-S4), suggesting the impaired β1AR response is not due to the loss of cGMP-PKG activity.

**Figure 1:**
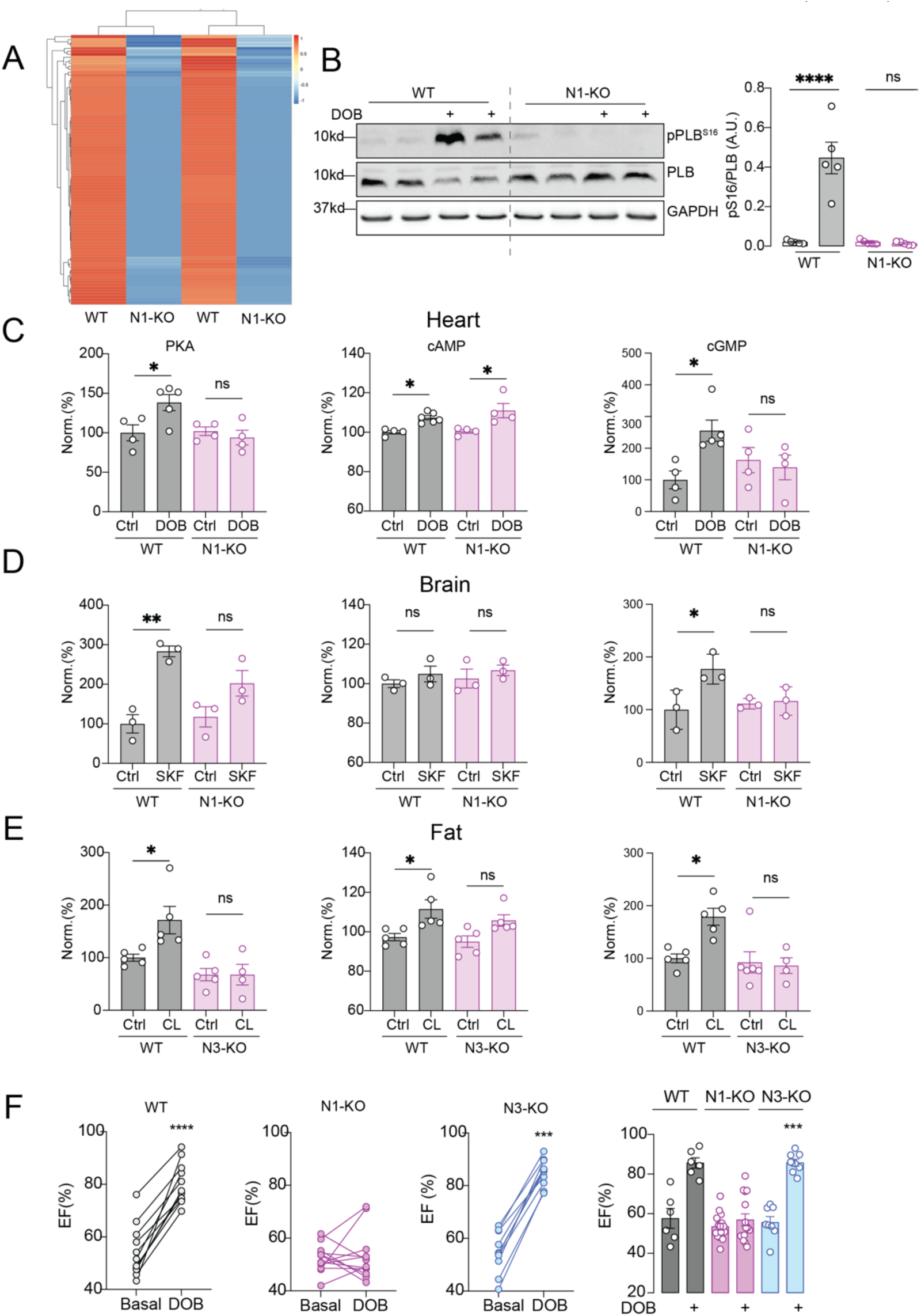
NOS is necessary for GPCR stimulation of PKA activity in heart, brain, and fat. (A) Heat map of proteins containing PKA-motif phosphosite peptides identified by mass spectroscopy after β₁-adrenergic receptor stimulation with dobutamine (D B, 100ng/g), showing altered phosphorylation of PKA target proteins in NOS1-KO hearts compared with WT controls. (B) Western blot of phospholamban (PLB) phosphorylation at Ser16 (pPLB16) induced by DOB in NOS1-KO hearts, with quantification. Data are presented as mean ± SEM, n=5, **** P < 0.0001 by One-way ANOVA followed by Tukey test. (C-E) ELISA assays of heart (C), brain (D), and brown adipose tissue (BAT) (D) lysates from WT and NOS1-KO or NOS3-KO mice after drug treatment with DOB, SKF81297 (SKF), or CL316,243 (CL) as indicated, for active PKA, cAMP, and cGMP. Data are presented as mean ± SEM, n = 3-5, *p < 0.05, **p < 0.01 by One-way ANOVA followed by Tukey test. (F) Echocardiography analysis of cardiac ejection fraction after injection of DOB in WT, NOS1-KO, and NOS3-KO mice, as indicated, plus summary. Data are presented as mean ± SEM, N = 8, ***P < 0.001, ****P < 0.0001 by Student’s paired *t*-test.

### GPCR signaling drives NOS isoform-dependent S-nitrosylation of PKA regulatory subunits across tissues

Building on these functional observations, we next explored interaction of β1AR with NOS1. We identified association of β1AR with NOS1 using co-immunoprecipitation (fig S5A). Additionally, using fluorescence co-immunostaining, the β1AR colocalized with NOS1 in WT but not NOS1-KO AVMs (fig S5B,C). The proximity of β1AR to NOS1 establishes a structural basis for NO production following β1AR stimulation. Because proximity to NOS has been associated with ability to be S-nitrosylated (28), we suspected that direct modulation of the activity of PKA by β1AR-NOS1-driven S-nitrosylation might be lost in NOS1-KO heart.

Analysis of S-nitrosylation using the biotin switch assay revealed that DOB stimulation induced S-nitrosylation of the PKA regulatory subunit RIIα in the heart, that ISO stimulation induced S-nitrosylation of both RIα and RIIα in the heart, that DOB or SKF stimulation induced S-nitrosylation of RIIβ in the brain, and that CL stimulation induced S-nitrosylation of RIIβ (but not PKA Cα subunit) in BAT (Figure 2A-C; fig S6A-D). This modification was NOS-dependent, as DOB-stimulated RIIα S-nitrosylation was abolished in NOS1-KO heart as were DOB- and SKF-stimulated RIIβ S-nitrosylation in NOS1-KO brain and CL-stimulated RIIβ S-nitrosylation in NOS3-KO BAT. We then employed FRET-based biosensors for cAMP (H187), cGMP (Gi500), and PKA activity (AKAR3) in isolated AVMs from wildtype and NOS-KO mice to assess the consequences of this S-nitrosylation. ISO stimulated cAMP, cGMP, and PKA activity to a similar extent in WT and NOS1-KO AVMs. However, while DOB increased cAMP comparably in WT and NOS1-KO cells, it failed to elevate cGMP level in the NOS1-KO AVMs. As expected, DOB-stimulated PKA activity also was reduced in the NOS1-KO cells (Figure 2D-G; fig S7). These findings suggest that NOS-dependent S-nitrosylation of PKA regulatory subunits may be necessary for GPCR activation of PKA, linking NO availability to the integrity of GPCR-cAMP-PKA signaling.

**Figure 2:**
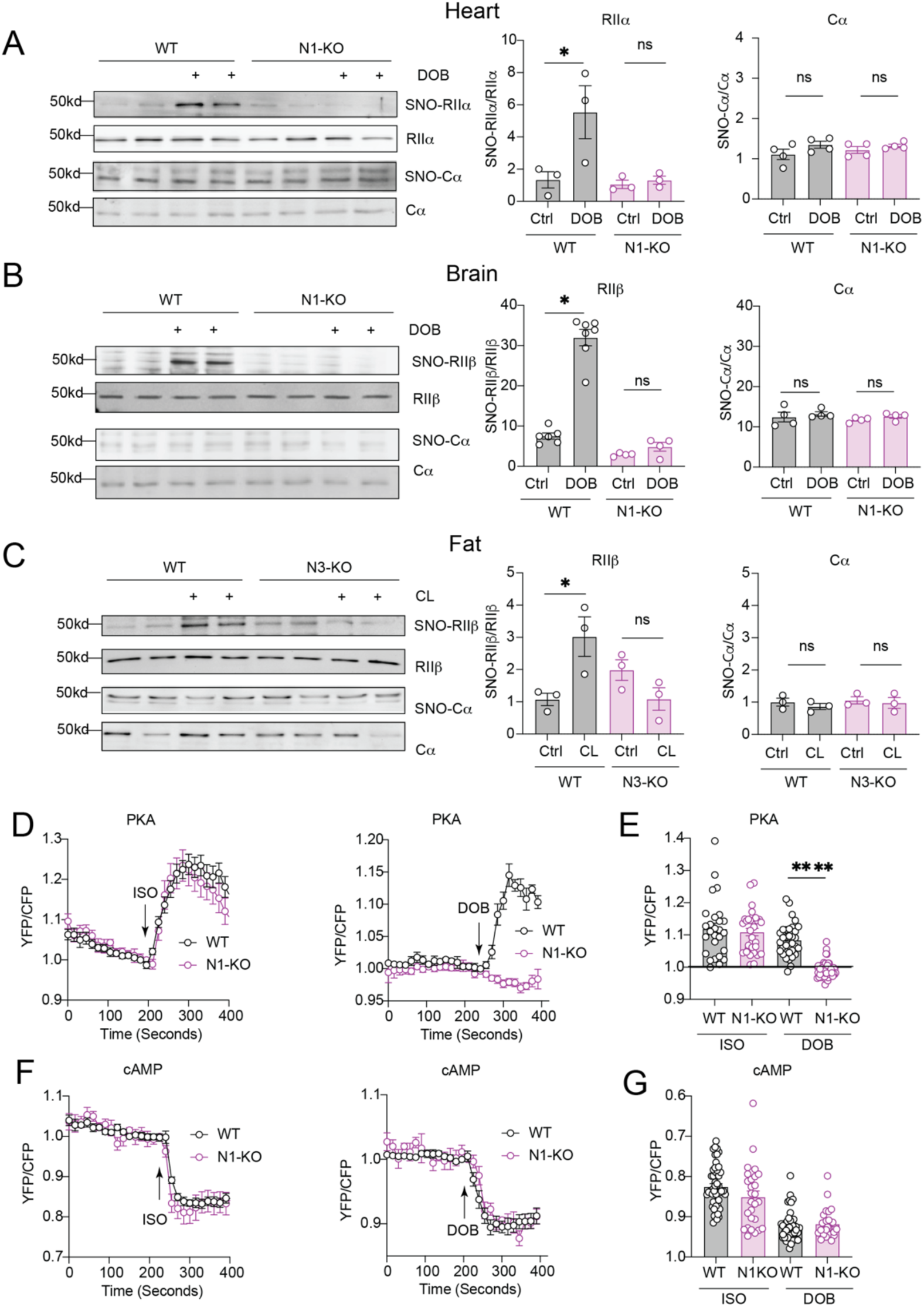
NOS is necessary for GPCR stimulation of S-nitrosylation of PKA regulatory subunits in heart, brain, and fat. (A) Detection of PKA RIIα S-nitrosylation by biotin-switch assay and Western blot in lysates from WT and NOS1-KO heart from mice treated with saline (control) or DOB. (B) Detection of PKA RIIβ S-nitrosylation by biotin-switch assay and Western blot in lysates from WT and NOS1-KO brain from mice treated with saline (control) or DOB. (C) Detection of PKA RIIβ S-nitrosylation by biotin-switch assay and Western blot in lysates from WT and NOS1-KO BAT from mice treated with saline (control) or CL. (*P < 0.05 by One-way ANOVA followed by Tukey test). (D, E) Isolated adult ventricular myocytes (AVMs) from WT or NOS1-KO mice expressing FRET biosensor for PKA activity (AKAR3) were treated with isoproterenol (ISO) or DOB. Data are presented as mean ± SEM. ****P< 0.0001 by 1-way ANOVA followed by Tukey test. (F, G) FRET biosensor for cAMP (H187) showing comparable ISO- and DOB-induced cAMP responses between WT and NOS1-KO AVMs. Data are presented as mean ± SEM.

### Mapping cysteine-dependent S-nitrosylation sites essential for holoenzyme activation

PKA RI and RII isoforms contain multiple cysteine residues. Two of these are conserved across all four RI and RII isoforms, while others are conserved only in RI or RII isoforms (fig S8A). Within PKA RIIβ and PKA RIIα, there are five conserved cysteines (Cys114, Cys147, Cys342, Cys371, Cys386) located within the two cAMP-binding domains. Comparing these residues across cAMP binding domains in other proteins as well as other nucleotide binding domains in signaling molecules reveals that these cysteine residues are found almost exclusively in the second cAMP binding domain of PKA RIIβ and PKA RIIα (Figure 3A; fig S9A, B). HEK293 cells overexpressing RIIβ variants with individual cysteine residues replaced by alanine mutations displayed reduced S-nitrosylation after treatment with the NO donor SNP, with greatest decrement in C147A, C371A and C386A (Figure 3B). Crystal structures of RIIβ holoenzymes show that the three cysteines (Cys342, Cys371, and Cys386), conserved in all RII subunits, are located in hydrophobic pockets in cyclic nucleotide binding domain B (CNB-B. fig S10A). Structural modeling predicts that nitrosylation of each cysteine would enhance the hydrophobicity of each site but would not sterically interfere with the fold, nor would it sterically clash with cAMP binding (fig S10A). Since no single mutation totally abolish nitrosylation, we generated a triple mutant (C342A/C371A/C386A) that abolished RIIβ S-nitrosylation in HEK293 cells treated with SNP (Figure 3D). Corresponding mutations were introduced at the homologous cysteine residues in RIIα (C327A/C356A/C371A), which similarly eliminated SNP-driven S-nitrosylation (Figure 3E). To determine whether cysteine S-nitrosylation affects the structure/function of PKA holoenzymes, we performed co-immunoprecipitation to probe for the binding between RIIβ and PKA catalytic subunit in HEK293 cells. The triple mutant RIIβ retained interaction with the catalytic subunit. While SNP enhanced dissociation of the catalytic subunit from WT RIIβ, this treatment did not release the catalytic subunit from the triple mutant (Figure 3F). We next assessed whether loss of these S-nitrosylatable cysteines impairs PKA activity. ELISA measurement of PKA catalytic activity in the immunoprecipitated complexes showed increased activity after SNP or ISO treatment in the WT condition, whereas no change was observed in the triple mutant (Figure 3G; fig S10B). To complement these biochemical findings, using HEK293 cells co-expressing the RII triple mutants with FRET biosensors. ISO dose response curves using cAMP FRET reporter showed preserved or even higher cAMP signals in the presence of either triple mutant RII isoform, in spite of markedly reduced PKA activity (Figure 3G; fig S10C). Taken together, these results demonstrate that the regulatory subunit’s S-nitrosylation facilitates the release of the catalytic subunit and is required for optimal PKA signaling.

**Figure 3:**
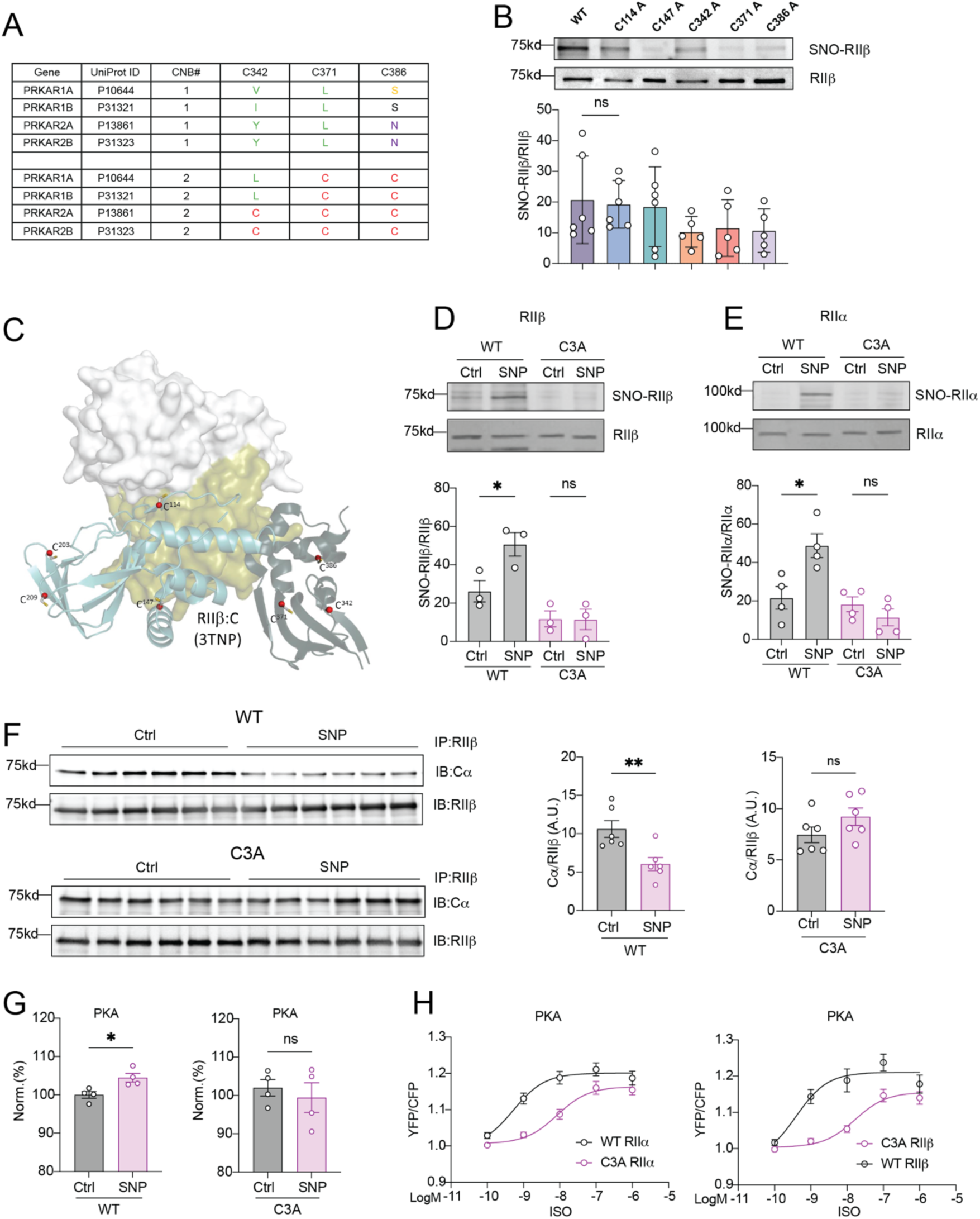
S-nitrosylation of PKA regulatory subunits. (A) Conserved cysteine residues (C342, C371, C386) in second cAMP-binding domain of PKA RIIβ and RIIα. Residue numbering according to RIIβ second cAMP-binding domain. (B) Identification of S-nitrosylated sites using biotin-switch assays in SNP-treated HEK293 cells overexpressing RIIβ variants with the indicated Cys to Ala mutations. Representative blot; group data are presented as mean ± SEM, n = 4-6. (C) In silico modeling localizing Cys342, Cys371, and Cys386 to the second cAMP-binding pocket of RIIβ subunit. (D) S-nitrosylation of WT versus triple mutant (C342A/C371A/C386A) RIIβ expressed in HEK293 cells and treated with SNP. (E) S-nitrosylation of WT versus triple mutant (C327A/C356A/C371A) RIIα in HEK293 cells and treated with SNP. Data in D,E are presented as mean ± SEM, n = 3-4, *P < 0.05 by One-way ANOVA followed by Tukey test. (F) Co-immunoprecipitation of overexpressed WT or triple-mutant RIIβ with PKA catalytic subunit in HEK293 cells treated with SNP. Data are presented as mean ± SEM, n = 6. **P < 0.01 by unpaired Student’s *t* test. (G) Co-immunoprecipitation of overexpressed PKA catalytic subunit with WT or triple-mutant RIIβ in HEK293 cells treated with SNP. PKA activity was assessed by ELISA in the immunoprecipitated complexes. Data are presented as mean ± SEM, n = 4. *P < 0.05 by unpaired Student’s t test. (H) PKA FRET AKAR3 biosensor was expressed with WT or mutant RIIα/β in HEK293 cells and stimulated with the indicated concentrations of ISO. ISO dose-response curves were analyzed by nonlinear regression, showing reduced PKA activity in cells expressing triple mutants. The log EC₅₀ values were −9.28 (WT) and −8.08 (RIIα mutant), and −9.41 (WT) and −7.78 (RIIβ mutant).

### GPCR-driven S-nitrosylation of PKA regulatory subunits via the SCAN nitrosylase

SCAN was identified recently as an enzyme that uses the S-nitroso group from S-nitroso-coenzyme A to place S-nitrosothiol onto specific sites on target proteins, and is encoded by the biliverdin reductase B (BLVRB) gene (27). We therefore investigated whether SCAN mediates the S-nitrosylation of PKA regulatory subunits following exposure to SNP and whether SCAN-mediated nitrosylation is necessary for PKA activation. SNP-induced nitrosylation of PKA RIIβ and RIIα was abolished in SCAN-KO HEK293 cells, demonstrating that SCAN is essential for regulatory subunit S-nitrosylation (Figure 4A). Using FRET biosensor assays, we compared cAMP levels and PKA activity in WT and SCAN-KO HEK293T cells upon GPCR activation with various ligands, including the non-selective β-adrenergic receptor agonist ISO (1 nM), the prostaglandin E2 receptor ligand PGE₂ (100 nM), the non-selective dopamine receptor D1-D5 agonist dopamine (10 µM), the GLP1 and GIP receptor dual-agonist tirzepatide (1 µM), the non-selective adenosine A1, A2A and A3 receptor agonist 5’-*N*-ethylcarboxamidoadenosine (NECA, 1 µM) or the dopamine D1 receptor agonist SKF-81297 (SKF, 10 µM). Although cAMP responses for all agonists were comparable or even higher in SCAN-KO cells than WT controls (Figure 4B), GPCR-stimulated PKA activity was reduced for all agonists (Figure 4C), indicating that SCAN-dependent S-nitrosylation is required for PKA activation downstream of cAMP. Dose-response curves with ISO, PGE₂, and NECA further confirmed these observations (Figure 4D–E; fig S11A-D), underscoring that cAMP elevation alone is not sufficient for full PKA activation and that SCAN-dependent S-nitrosylation provides the additional requirement. Mirroring the PKA activity dysfunction observed with the FRET-based biosensors, ISO (1 nM) stimulation-induced robust PKA-substrate phosphorylation in WT but not in SCAN-KO cells (Figure 4F; fig S11E). Our results identify SCAN as the enzyme mediating GPCR-stimulated S-nitrosylation of PKA regulatory subunit s-nitrosylation, revealing this previously unrecognized requirement for full GPCR signaling.

**Figure 4:**
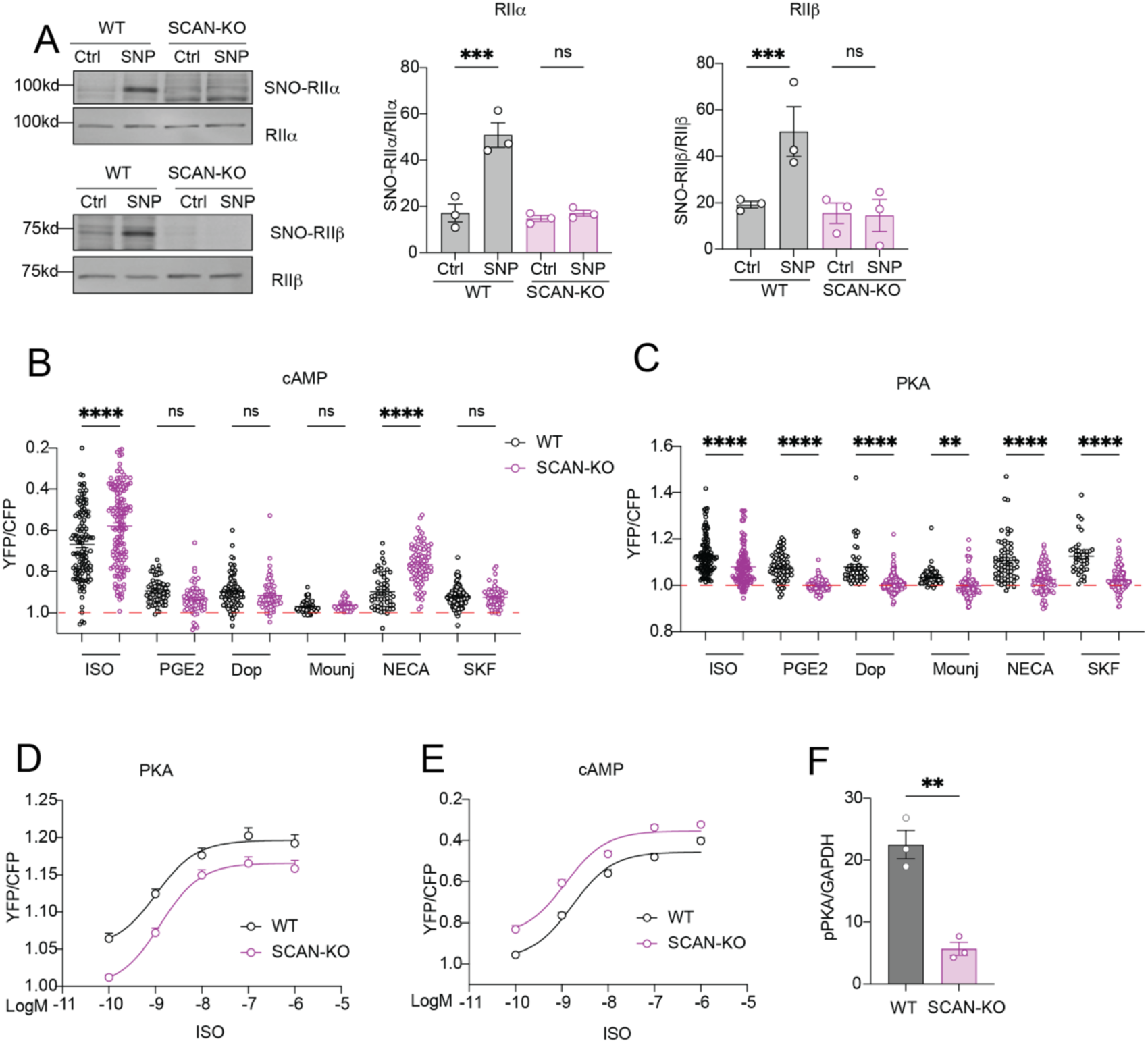
SCAN mediates S-nitrosylation of PKA regulatory subunits to regulate PKA activation after GPCR stimulation. (A) SNP-mediated S-nitrosylation of PKA RIIβ or RIIα, detected by western blot of biotin-switch assays of SNP-treated WT and BKVRB-KO HEK293T cells overexpressing PKA RIIβ or RIIα. ***P < 0.001 by One-way ANOVA followed by Tukey test. (B, C) Plasma-membrane–targeted FRET biosensors for PKA activity (panel B, PM AKAR3) and cAMP (panel C, PM H187) and in WT and SCAN-KO HEK293T cells stimulated with ISO (1 nM), PGE₂ (100 nM), dopamine (Dop, 10 µM), tirzepatide (tirz, 1 µM), NECA (1 µM), or SKF-81297 (SKF, 10 µM).**P < 0.01, ****P < 0.0001 was determined by one-way ANOVA with Tukey’s multiple comparison test. (D, E) Dose–response curves for ISO-stimulated PKA by FRET (D) and cAMP by FRET (E), generated by nonlinear regression analysis showing preserved or elevated cAMP responses in SCAN-KO cells (log EC₅₀ = −8.91 for WT, −8.97 for SCAN-KO) but consistently reduced PKA activity (log EC₅₀ = −8.98 for WT, −8.90 for SCAN-KO). **P < 0.01, ****P < 0.0001 was determined by one-way ANOVA with Tukey’s multiple comparison test. (F) ISO (1 nM)-induced PKA phosphorylation of substrates quantified in WT and SCAN-KO cells, from anti-phospho-PKA substrate (RRXS*/T*) blots in Figure S10E. Data are presented as mean ± SEM from n = 3-5, **P < 0.01 by Student’s *t* test.

### NOS1-derived NO is required for β1-adrenergic receptor stimulation of PKA activity and cardiac contractility

Given the impaired β1AR-PKA signaling in the absence of NOS1, we investigated whether restoring NO availability could rescue full PKA activation. FRET-based biosensors were expressed in isolated AVMs from WT and NOS1-KO mice. DOB stimulation resulted in comparable increases in cAMP in NOS1-KO or after SNP treatment, indicating that neither NOS1 deletion nor SNP supplementation altered cAMP generation (Figure 5A). As controls, cGMP levels were enhanced in WT cells but blunted in NOS1-KO AVMs, and pretreatment with SNP fully restored cGMP levels to those of WT cells (Figure 5B). Moreover, PKA activation was attenuated after DOB stimulation in NOS1-KO cells, but this was rescued by SNP pretreatment; pretreatment with SNP fully restored PKA levels to those of WT cells (Figure 5C). Consistent with these FRET signaling results, sarcomere shortening was used as an *in vitro* index of contractility in isolated AVMs. In WT cells, DOB stimulation enhanced sarcomere shortening relative to baseline, whereas SNP alone had minimal effect, while combining SNP and DOB resulted in a further, non-significant increase (Figures 5D,E). In NOS1-KO AVMs, DOB or SNP alone failed to augment sarcomere shortening, but SNP followed by DOB increased contractile responses to levels comparable to those of WT (Figures 5F,G). In agreement. DOB-induced PLB phosphorylation was attenuated in the NOS1-KO cells relative to WT cells. However, pretreatment with SNP followed by DOB restored PLB phosphorylation in NOS1-KO cells (Figure 5H). To establish the *in vivo* relevance of these cellular findings, we assessed the cardiac function using echocardiography in WT and NOS1-KO mice. In WT mice, DOB injection enhanced EF compared with baseline, whereas this response was markedly blunted in NOS1-KO mice. Pretreatment with SNP rescued the attenuated DOB response in the KO mice, restoring EF to a level comparable to that of WT mice (Figure 5I; table S5). These results underscore that NOS1-derived NO is essential for β1AR-induced PKA activation, and cAMP alone is not sufficient for full PKA activity in heart.

**Figure 5:**
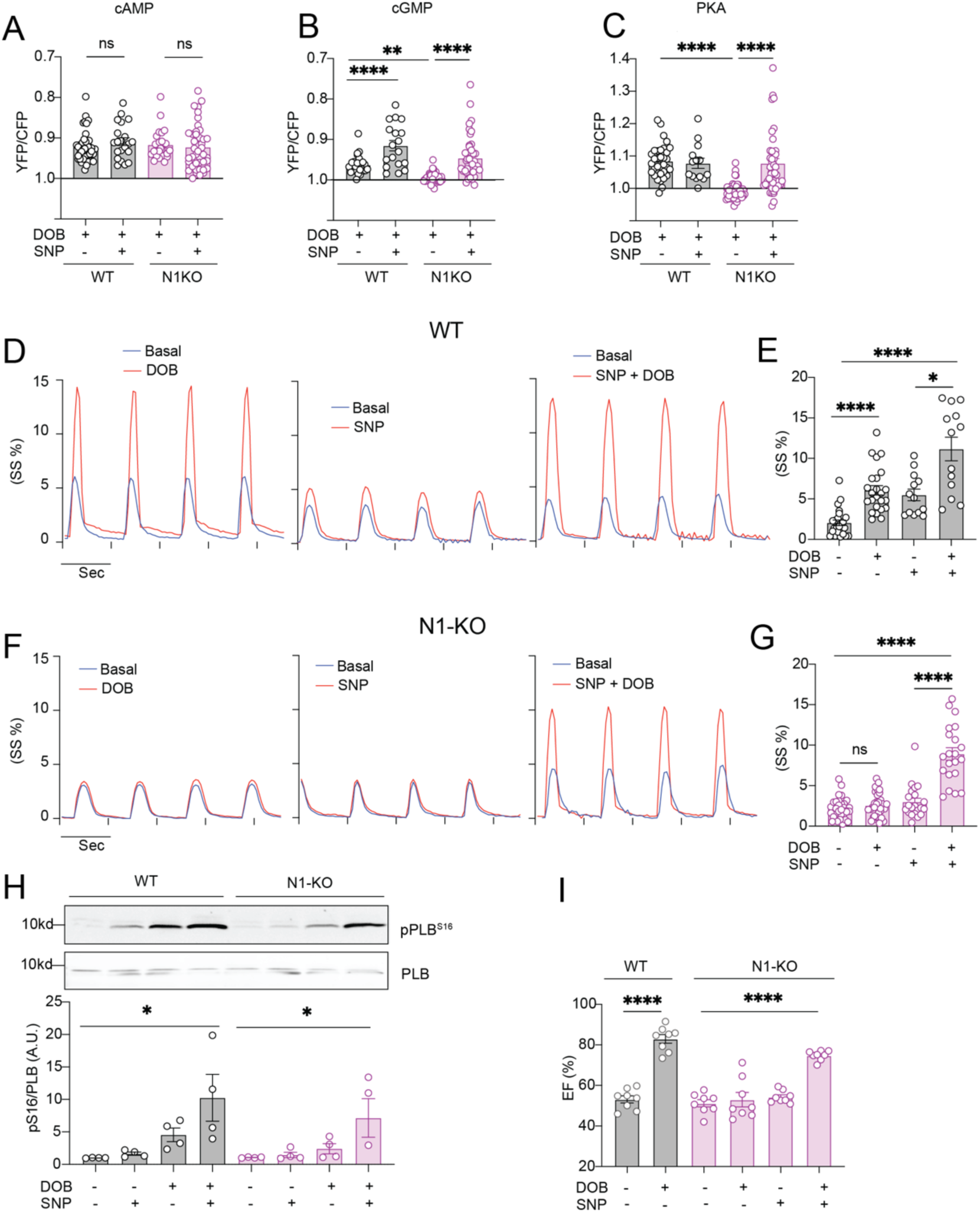
Clinical NO donor drug rescues β-adrenergic responsiveness in NOS1 KO hearts. (A–C) FRET biosensors for cAMP (panel A, H187), cGMP (panel B, Gi500), and PKA activity (panel C, AKAR3) in isolated AVMs from WT and NOS1-KO mice before and after treatment with DOB or SNP as indicated. (D, E) WT AVMs were paced at 1Hz, and sarcomere shortening (SS) was measured at baseline and after treatment with DOB or SNP as indicated. Representative traces (D) and quantification of sarcomere shortening (SS%) in WT AVMs (E). (F, G) NOS1-KO AVMs paced at 1Hz, and sarcomere shortening was measured at baseline and after treatment with DOB or SNP as indicated. Representative traces (F) and quantification of sarcomere shortening (SS%) in NOS1-KO AVMs (G). (H) Immunoblot analysis of PLB phosphorylation at Ser16 (pPLB16) after treatment with DOB and SNP as indicated, with quantification below. Data are presented as mean ± SEM; 3-4 independent experiments per condition. (I) Ejection fraction measured by echocardiography in WT and NOS1-KO mice at baseline and after treatment with DOB or SNP as indicated. Data are from 8-10 mice. \**P* < 0.05, \*\**P* < 0.01, \*\*\**P* < 0.001, ****P< 0.0001 by one-way ANOVA with Tukey’s multiple comparison test.

### Disrupted β1-adrenergic receptor-NOS1 axis in failing hearts impairing cAMP-PKA signaling is restored by NO repletion

We established a myocardial infarction (MI) mouse model by ligating the left anterior descending coronary artery to investigate how heart failure remodels the β1AR-NOS1 signaling pathway. Western blot analysis showed marked reductions in NOS1, β1AR, and the anchoring protein SAP97, whereas NOS3 expression was increased, indicating a compensatory adaptations (Figure 6A,B). To explore translational relevance, analysis of human cardiac mRNA sequencing data from healthy controls and patients with heart failure with reduced ejection fraction revealed reduced SCAN transcript levels, elevated NOS3 expression, with no significant change in NOS1 (fig S12A-C). Co-immunoprecipitation further demonstrated diminished interaction of NOS1 with β1AR in MI hearts (Figure 6C,D). In line with these molecular changes, ELISA assays revealed reduced PKA activity in MI hearts, despite preserved DOB-induced cAMP generation (Figure 6E,F). Given the blunted PKA activity in MI hearts, we analyzed phosphorylation of PLB at Ser16. In MI hearts, DOB treatment failed to significantly increase PLB phosphorylation compared with saline, whereas SNP pretreatment followed by DOB markedly enhanced phosphorylation, yielding a robust and significant elevation compared with the saline group (Figure 6G). Consistent with these biochemical findings, echocardiography showed that in sham mice, DOB stimulation increased EF, with no further effect of SNP pretreatment (Figure 6H). In contrast, EF was unchanged after DOB or SNP alone in MI hearts, but SNP pretreatment followed by DOB significantly increased EF in MI mice (Figure 6H; table S6). Our data demonstrate that MI disrupts the β1AR-NOS1 axis, uncoupling cAMP generation from PKA activity, and that NO availability is necessary to restore β1AR-dependent contractile responses in the failing heart, whereas exogenous NO donor (SNP) restores PKA S-nitrosylation and rescues β₁AR-mediated contractile function (Figure 6I).

**Figure 6:**
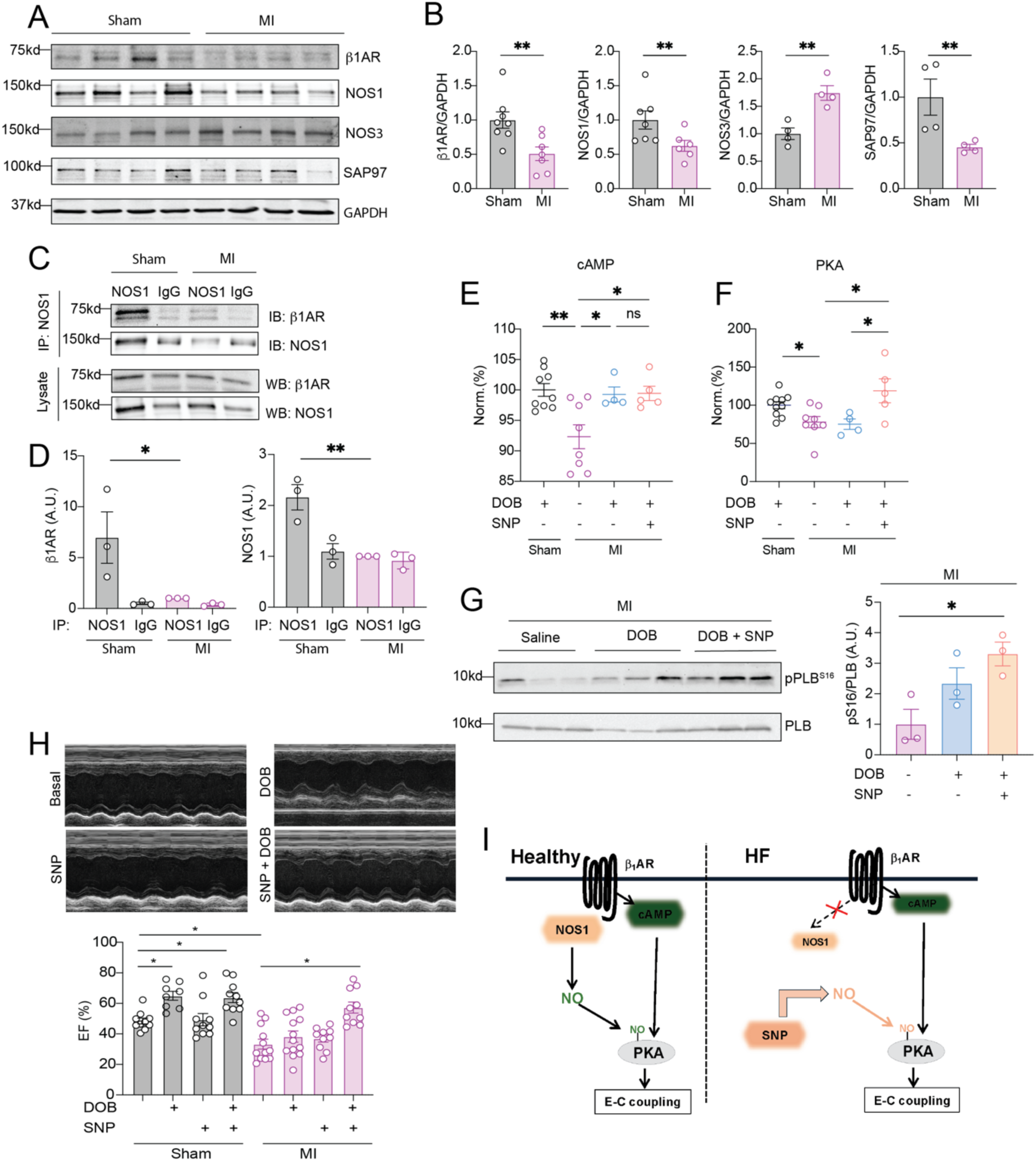
Clinical NO donor drug rescues DOB stimulation of cardiac ejection fraction in heart failure. (A, B) Western blot detection (A) of β1-adrenergic receptor, NOS1, NOS3, and SAP97 protein expression in hearts of WT mice with Sham surgery or LAD ligation (MI), with quantification (B). **P < 0.01 by Student’s unpaired *t*-test. (C, D) Co-immunoprecipitation of NOS1 with β1AR in MI hearts (C), with quantification (D). N = 3 **P < 0.01 by Student’s unpaired *t*-test. (E, F) cAMP (E) and PKA activity (F) measured by ELISA in Sham and MI hearts pretreated with DOB or SNP as indicated. (G) Western blot analysis of phospholamban phosphorylation at Ser16 (pPLB16) in heart from DOB- and SNP-treated MI mice. (H) Echocardiography measurement of cardiac ejection fraction after injection of DOB and SNP as indicated in Sham and MI mice. Data are presented as mean ± SEM, and \**P* < 0.05, \*\**P* < 0.01 by one-way ANOVA with Tukey’s multiple-comparison test.

## Discussion

Activation of many GPCRs subtypes leads to increases in cAMP and in NO, two critical secondary messengers for physiological responses throughout the body (29,30). While cAMP predominantly activates PKA for a broad range of physiological responses, such as neuronal excitation, muscle contraction, hormone secretion, and fuel metabolism, NO is primarily known for promoting vessel relaxation via PKG (31–35). Here, we demonstrate that NO is required for PKA activation. NO directly modifies PKA regulatory subunits via S-nitrosylation after stimulation of multiple GPCR types in brain, heart and fat tissues. This S-nitrosylation is mediated enzymatically by SCAN, occurs on conserved cysteine residues, and promotes dissociation and activation of PKA holoenzymes to promote PKA kinase activity. Deleting NOS1 or NOS3 blunts GPCR-stimulated PKA activation, despite normal increases in cAMP. Moreover, in the transgenic NOS1-deleted heart and in a heart failure model with reduced NOS1 expression, treatment with the NO donor drug nitroprusside successfully restored β1-adrenergic stimulation of PKA activity, substrate phosphorylation, myocyte shortening, and cardiac contractility. Together, our data define an unexpected paradigm in which GPCR stimulation of NO production is required for cAMP-mediated PKA activation, overturning the prevailing model of GPCR transduction. Our data also suggest important new uses for the drug sodium nitroprusside in the clinical setting.

PKA holoenzyme is a tetramer consisting of two R subunits and two C subunits, in which the R subunits serve to inhibit the catalytic activity of C subunits (36,37) but also to localize them to sites of action (38). In the classic paradigm, stimulation of GPCRs leads to increased cAMP levels (3,39). The cAMP then binds to the two cAMP binding domains in R subunits (40,41) to promote dissociation of catalytic subunits, freeing them to phosphorylate substrate proteins (42). Our data show that S-nitrosylation occurs at conserved cysteine residues (Cys342, Cys371, and Cys386 in RIIβ) that are localized exclusively in the second cAMP binding domain. Mutation of these cysteine residues impairs PKA activation (despite elevated cAMP induced by GPCR ligands) by blocking dissociation of the PKA holoenzyme and prevents PKA activation by SNP. Structural analysis predicts the S-nitrosylation of Cys342, Cys371, and Cys386 would enhance the hydrophobicity of the cAMP binding pocket, which would lock the R-subunit in a more stable cAMP-bound conformation. In contrast, mutating Cys to Ala would reduce hydrophobicity, making the Cys mutant harder to be activated with cAMP. A previous report found that S-nitrosylation of the PKA RIα subunit by NO caused release of active PKA catalytic subunits in the apparent absence of cAMP (43), but native regulation by GPCRs and site of S-nitrosylation were not defined. In addition, several SNO-proteomics studies have identified PKA-RI and PKA-RII as endogenous SNO-proteins in various tissues and under various conditions, but these have never been validated (44–46). Our work should motivate further studies to understand the structural basis of S-nitrosylation effects on PKA and its physiological implications across tissues.

GPCR activation of NO signaling, and particular its signaling via S-nitrosylation, have been broadly underappreciated, despite nearly all GPCR signaling components having been reported as S-nitroso-proteins in proteomic screens (24) implying a broad role in signal transduction. Here we provide a clear example of integration of GPCR–NO–S-nitrosothiol signaling and canonical GPCR–Gs protein–cAMP signaling, defining PKA as a conditional effector requiring convergence of both signals and raising the question of the extent to which GPCR-driven NO signaling attributed to PKG is actually mediated by S-nitrosylated PKA. The traditional downstream effector of NO is sGC, which produces cGMP to activate PKG (47,48). While this pathway is well known for relaxation of smooth and cardiac muscle cells, their roles in other tissues are less defined as PKA and PKG share many substrates involved in physiological response (49,50). Interestingly, sGC has been reported to act as a transnitrosylase enzyme to transfer S-nitrosothiol from itself to other proteins including Rho kinase, and mutation of the sGC S-nitrosylation site leads to defective endothelial cell vasorelaxation responses and elevated blood pressure (51). Additionally, cGMP can crosstalk with cAMP via modulation of dual-specificity PDE2 and PDE3, which hydrolyze both cAMP and cGMP (52,53). Our discovery of S-nitrosylation as a direct regulator of PKA activation identifies a new aspect of this sophisticated regulatory system, offering a simple and universal mechanism to regulate cyclic nucleotide-dependent protein kinase stimulation by various neurohormones.

Deficiency in NOS1 and NOS3 expression and activity has been implicated in a broad range of diseases in brain, heart, fat, liver, lung, kidney, muscle, and vessels (54–58). Further, inducible NOS (NOS2) is often elevated in chronic diseases and inflammation, and produces pathological levels of NO and other reactive species (59,60). Despite extensive efforts, directly targeting NOS or NO has not yet been successful in drug development and therapy (61–64). Our studies suggest reinterpretation of results in terms of enzymatic S-nitrosylation by direct analogy to phosphorylation, and reveal a critical role for GPCR-mediated S-nitrosylation in cellular signaling that has been overlooked in development of current therapeutics. Our data suggest that the coupling of β1AR to NOS1 is disrupted in heart failure, indicating that restoring the coupling between GPCRs and NOS enzymes may be an alternative approach to rescuing the NO signal in diseases. Moreover, our work points to a new clinical application for SNP and reevaluation of its role in heart failure. Heretofore, SNP has been used to reduce afterload (blood pressure) in heart failure settings. It was not appreciated that SNP may directly improve contractility through PKA activation or that its inotropic effects would depend on SNO-status. Thus, our data suggest a more nuanced approach to heart failure treatment, including potential uses for NO-based therapies.

Finally, we have revealed that the nitrosylase enzyme SCAN is critical for GPCR-mediated S-nitrosylation and activation of PKA. Deleting SCAN blunts PKA activation by multiple GPCR ligands, including isoproterenol, adenosine, dopamine, prostaglandin E2, and the clinical drug tirzepatide. Deletion of SCAN also leads to increased insulin-induced Akt activation due to failure to S-nitrosylate the insulin receptor and IRS-1 (12,27). These studies present an emerging paradigm of NO modulation of critical kinases and signaling molecules in physiology, which is likely to be widespread among GPCR signaling effectors and mediators as well (24). Enzymatic crosstalk among kinases and nitrosylases downstream of receptor activation may represent a new field of study.

In summary, we have identified S-nitrosylation of PKA R subunits as a common mechanism critical for activation of PKA downstream of GPCR stimulation in multiple tissues, acting in concert with cAMP. These results change our understanding of GPCR-induced PKA regulation—and of GPCR transduction more generally and suggest new therapeutic opportunities and treatment strategies in a broad range of diseases.

## Methods

### Animals

Wild-type (WT) C57BL/6J (Stock #000664), β1-adrenergic receptor knockout (β1AR-KO), neuronal nitric oxide synthase knockout (NOS1-KO), endothelial nitric oxide synthase knockout (NOS3-KO), and Protein kinase G type II knockout (PKG2-KO) mice were obtained from The Jackson Laboratory (Bar Harbor, ME, USA). Protein kinase G type I-flox (PKG1-F/F) was described previously (65) and crossed with MHC-cre (IMSR_JAX:009074) to generate PKG1-conditional KO (PKG1 cKO) as well as PKG1-cKO/PKG2-KO. All genetically modified mouse lines were backcrossed onto the C57BL/6J background, and C57BL/6J littermates or strain-matched C57BL/6J mice were used as controls. WT, NOS1-KO, and NOS3-KO groups included only male mice, while the β1AR-KO, PKG1-F/F, PKG1-cKO, and PKG2-KO groups included both males and females; however, sex was not considered a variable in the analysis. Mice were 2–3 months old at the time of the experiments. All animals were housed in a facility accredited by AAALAC under standard conditions (22 ± 1 °C, 40–60% humidity, 12-hour light/dark cycle with lights on at 07:00) with ad libitum access to standard chow and water. They were group-housed (3–5 per cage) in ventilated cages with corn cob bedding. Animals received at least one week of acclimatization after arrival before testing. All procedures were conducted in accordance with the NIH Guide for the Care and Use of Laboratory Animals and received approval from the Institutional Animal Care and Use Committee at the University of California, Davis (protocol #23640). Mice were randomly assigned to experimental groups.

Heart tissue from the left ventricle and brain tissue were obtained from WT and NOS1 KO mice. Brain slices were treated with either saline or DOB at 100 nM, while the heart was Langendorff perfused with the same solutions for 10 minutes. Brain tissues were also harvested WT and NOS1 KO mice received an intraperitoneal injection of either saline or the dopamine D1 receptor agonist SKF-81297 hydrobromide (0.5 mg/kg) for 10 minutes. Interscapular brown adipose tissue (iBAT) was isolated from WT and NOS3 KO mice after injection with either saline or CL 316,243 (10 μg/kg) for the same duration as DOB. Additionally, we used two other mouse models: MI mice injected with DOB (400 µg/kg, i.p.) and pretreated with SNP (100 µg/kg), followed by the same dose of DOB. All tissues were collected rapidly, flash frozen in liquid nitrogen, and stored at −80 °C for S-Nitrosylation detection, TMT phosphoproteomic, ELISA, and Western blot assays.

### Myocardial Infarction (MI)

Male mice aged two to three months were used to induce myocardial infarction (MI) through permanent ligation of the left anterior descending (LAD) coronary artery. Mice were given 100% oxygen (0.8 L/min), anesthetized with isoflurane (5% for induction and 2-3% for maintenance), and ventilated with a rodent respirator at 110–130 breaths/min with a tidal volume of 0.2 mL. A left thoracotomy was done to reveal the heart following endotracheal intubation, and the LAD was observed under a surgical microscope. An 8-0 suture was used to ligate the artery, resulting in a left ventricular infarction of about 30–40%. Electrocardiography confirmed the infarction’s onset. Sham-operated mice underwent the same surgical procedure, including thoracotomy and cardiac exposure, but without LAD ligation. Postoperative analgesia was administered with buprenorphine (0.1 mg/kg, subcutaneously) immediately after surgery and during recovery. Mice were closely monitored until fully awake and then housed under standard conditions. Cardiac function was assessed by transthoracic echocardiography to confirm the establishment of MI.

### Transthoracic Echocardiography

Cardiac function was evaluated in WT, NOS1-KO, NOS3-KO, β1AR-KO, PKG1-F/F, PKG1-cKO, PKG2-KO, and PKG1-cKO/PKG2-KO mice using transthoracic echocardiography using the Vevo 2100 Imaging System and a 22- to 55-MHz linear probe (Visual Sonics). Mice were lightly anesthetized with isoflurane (1–2% inhalation in 100% O₂) and placed supine on a heated platform maintained at 37 °C, with continuous ECG and respiratory monitoring. One day before imaging, chest hair was removed to improve acoustic coupling. Two-dimensional (2D) short-axis images at the mid-papillary level were captured in M-mode to measure multiple parameters, such as interventricular septal thickness, left ventricular internal diameter, posterior wall thickness, ejection fraction, fractional shortening, left ventricular mass, and volume. All parameters were assessed during systole and diastole. Baseline data were collected after stabilization, followed by intraperitoneal injection of DOB (400 µg/kg), ISO (100 µg/kg), or SNP (100 µg/kg) in sterile saline. Ten minutes post-injection, echocardiographic measurements were repeated to evaluate the drug’s peak effect. All data were analyzed offline by an investigator blinded to genotype and treatment condition using Vevo LAB analysis software.

### Myocyte isolation

AVMs were isolated from 8- to 12-week-old WT and NOS1-KO mice, following the previously reported protocol (66). Hearts were cannulated to a Langendorff apparatus and were first perfused through aorta canulation with an isolation buffer (in mmol/L): NaCl 120, KCl 5.4, NaH2PO4 1.2, NaHCO3 20, MgSO4 1.2, Glucose 5.6, 2,3-Butanedione monoxime 10, Taurine 20; PH 7.33. The heart underwent another perfusion round with a solution of isolation buffer containing 0.05% collagenase type II and 0.01% protease type XIV, without calcium. This was followed by flushing with isolation buffer containing 0.05% collagenase type II, 0.01% protease type XIV, and 50 μmol/L CaCl2. The flushed hearts were cut below the atria into a plastic container filled with 5 mL of digestion buffer. They were gently triturated before being transferred into the stopping buffer (isolation buffer with 10% fetal bovine serum, FBS). Isolated AVMs were used immediately for excitation-contraction Coupling and western blot and after 36 hours for FRET assays and western blot.

### Cell Culture

HEK293 QBI cells were maintained in Dulbecco’s modified Eagle’s medium (DMEM, Gibco) supplemented with 10% fetal bovine serum (FBS, Gibco) and 1% penicillin–streptomycin (Gibco) at 37 °C in a humidified 5% CO₂ incubator. Cells were subcultured at approximately 70–80% confluency using trypsin and seeded according to the needs of each experiment. Transient transfections were performed using Lipofectamine 3000 (Thermo Fisher Scientific, L3000015) according to the manufacturer’s protocol. To minimize serum interference, cultures were switched to serum-free DMEM 4 h before transfection, and the medium was refreshed 6-8 h later. For PKA regulatory subunit experiments, RIIβ constructs were expressed with an N-terminal FLAG tag and mCherry, whereas RIIα constructs carried a FLAG tag and tdTomato. These reporters allowed visual confirmation of expression before use and supported FLAG-based immunoprecipitation in downstream assays.

HEK293T WT and SCAN/biliverdin reductase B knockout (SCAN KO) cells were described previously (27) and were maintained under identical growth conditions to the QPI line. Culture, passaging, and transfection procedures were performed as described above unless otherwise noted. For S-nitrosylation experiments, HEK293 cells were transfected with wild-type (WT) or cysteine-to-alanine mutant constructs of PKA regulatory subunits. RIIβ mutants included C114A, C147A, C342A, C371A, C386A, and a triple mutant (C342A/C371A/C386A). RIIα mutants included a combined C271A/C327A/C356A mutant. All constructs carried FLAG tags, and fluorescent reporters were included as described above to aid expression verification. Thirty-six hours after transfection, cells were treated with sodium nitroprusside (SNP, 200 µM, 7 min, 37 °C) prior to S-nitrosylation detection by biotin switch assay.

### Fluorescence resonance energy transfer (FRET)

Isolated AVMs were plated on laminin-coated coverslips and cultured in M1018-based FRET media, including 10 mM HEPES, 6.25μM blebbistatin, 0.2% Bovine Serum Albumin (BSA), pH 7.35, and 10% FBS) for 2 h, then the media was switched to serum-free (SF) M1018-based FRET media containing FRET-based biosensor for cAMP (H187), FRET-based biosensor for cGMP (Gi500), FRET-based biosensor for PKA activity (AKAR3) were expressed in AVMs by infection with recombinant adenovirus (MOI 100) for 36 hrs before measuring the FRET responses to isoproterenol, and dobutamine stimulation (67,68). The FRET-based biosensor H187 (Epac-SH187), Gi500, or AKAR3 (69) were generated using the AdEasy system (Qbiogene) (70).

Wild-type (WT) HEK T cells and HEK T SCAN knockout (KO) cells were cultured under the same conditions. Cells were seeded on coverslips in 24-well plates at roughly 60–80% confluency about a day before transfection with either AKAR3 or H187 biosensors (pcDNA3.1 constructs) using the same Lipofectamine™ 3000 protocol, with a total of 500 ng plasmid DNA per well in 24-well plates. We also co-transfected WT HEK293 cells with pcDNA3.1 constructs that encode AKAR3 or Epac-SH187 (71,72), along with wild-type or mutant PKA regulatory subunits (RIIβ or RIIα; 1:1 ratio of biosensor and PKA subunit constructs). The RIIβ mutant carried three cysteine-to-alanine substitutions (C342A, C371A, C386A), whereas the RIIα mutant contained analogous substitutions at C271A, C327A, and C356A. After 15 minutes of complex formation at room temperature, the transfection mixture was added to the cells. Cells were returned to incubation for 18-30 h, after which they were used for FRET imaging. For plasma membrane– targeted biosensors, cells were infected with recombinant adenoviruses encoding PM-H187 or PM-AKAR3 (73) at an MOI of 100 for 12 h in serum-free medium before imaging.

Cells were rinsed and kept in calcium-free PBS. Images were acquired using a Hamamatsu Orca-Flash 4.0 camera (Bridgewater, NJ) managed by Metafluor software on the A1 inverted Zeiss AXIO Observer fluorescence microscope. Donor fluorophore (cyan fluorescent protein, CFP) was excited at 430-455 nm; The images were obtained using an emission fluorescent filter with a wavelength of 475DF40 (for CFP) and 535DF25 (for a yellow fluorescent protein, YFP). The interval of image acquisition was 20 seconds, and the exposure time was 200ms per channel. The images from both channels underwent background subtraction, and the ratio of yellow to cyan color was computed at various time intervals. When cAMP binds to H187, it leads to a decrease in the YFP/CFP ratio (74).

In FRET assays, AVMs were exposed to DOB (100 nM, Sigma-Aldrich, D0676) or ISO (10 nM, Sigma-Aldrich, I6504), and SNP (200 μM, Sigma-Aldrich, 13755-38-9). HEK cells were exposed to ISO with a wider concentration range (100 pM to 1 μM). Other GPCR agonists included prostaglandin E₂ (1 nM to 10 μM), SKF-81297 hydrobromide at 10 μM (Tocris Bioscience, Bristol, UK; 1447), NECA (1 nM to 10 μM, MedChemExpress; HY-13465, Monmouth Junction, NJ, USA), tirzepatide acetate (1 μM, MedChemExpress, HY-134260A), and dopamine hydrochloride (10 μM, Sigma-Aldrich, H8502).

### Excitation-Contraction Coupling in Isolated Adult Ventricular Myocytes

Freshly isolated AVMs were recovered for 10 min in beating buffer (in mM: 120 NaCl, 5.4 KCl, 1.2 MgSO4, 1.2 NaH₂PO₄, 20 HEPES, 5.5 glucose, 1.0 CaCl₂; pH adjusted to 7.1 with NaOH) at room temperature. For imaging, a glass-bottom dish containing 3 mL of beating buffer was placed on the stage of a Zeiss AX10 inverted fluorescence microscope (Carl Zeiss, Dublin, CA, USA). We added the dye-loaded myocytes gently to the center of the dish. After that, Platinum electrodes connected to an SD9 stimulator (Grass Technologies, Warwick, RI) were immersed in the bath near the cells. The stimulator delivered field pulses at 1 Hz with a 20-ms delay, 20-ms pulse width, and 50-V output. Image acquisition was performed using Metamorph software, capturing 5 consecutive contractions (25 ms exposure, 200 frames) under baseline conditions and 5 min after in vitro drug application, as indicated in each experiment. Where applicable, cells were pretreated for 7 min with SNP at 200 μM prior to recordings. Sarcomere shortening (SS) was calculated from bright-field recordings as: FS (%) = (maximal cell length − minimal cell length) / maximal cell length × 100. For statistical analysis, one dish per condition was prepared for each animal, and data from at least three animals were included per experimental group.

### S-Nitrosylation Detection (Biotin Switch Assay)

S-nitrosylated proteins were detected using a commercial detection Kit (Cayman Chemical) according to the kit protocol. We washed the cells three times with ice-cold buffer, followed by scraping. Cells were pelleted by centrifugation for 5 minutes at 500 × g at 4 °C and resuspended in buffer A containing blocking buffer. The resulting lysates were clarified by another spin (10 minutes, 4 °C), and proteins were precipitated by adding four volumes of ice-cold acetone and leaving the mixture at –20 °C for 1 h. The pellets were then dissolved in buffer B with the reducing and labeling reagents and kept for 1 h at room temperature in the dark. Following this labeling step, we used cold acetone to precipitate the proteins. The final pellets were resuspended in wash buffer and processed for immunoprecipitation. After biotin-switch assay and immunoprecipitation, the protein samples were loaded onto SDS-PAGE gels for separation by size. Once resolved, they were transferred to PVDF membranes to create a stable platform for downstream probing with antibodies. To minimize nonspecific binding, membranes were blocked in 2% BSA prepared in TBS (pH 7.4) for 1 h at room temperature, followed by incubation with S-nitrosylation (biotin) detection reagent I (HRP) diluted 1:75 in 2% BSA for 1 h. After washing, proteins were visualized using ECL and imaged with a ChemiDoc system (Bio-Rad). For confirmation of target identity, we re-probed the membranes with FLAG antibody or other target-specific primary antibodies overnight at 4 °C, followed by HRP-conjugated secondary antibodies. Band intensity was quantified using Image Lab software.

### ELISA of PKA Activity and cAMP and cGMP Quantification

PKA kinase activity in tissue lysates was measured using a commercial kit (Abcam, ab139435) following the manufacturer’s protocol. Briefly, tissue was lysed in buffer containing 20 mM MOPS, 50 mM β-glycerophosphate, 50 mM sodium fluoride, 1 mM sodium orthovanadate, 5 mM EGTA, 2 mM EDTA, 1% NP-40, 1 mM DTT, 1 mM benzamidine, 1 mM PMSF, and 10 µg/mL leupeptin and aprotinin. Afterward, BCA analysis provided protein concentration. Equal protein amounts (30 µL) were added to substrate-coated wells, and the addition of 10 μl ATP initiated the kinase reaction. Following incubation at 30 °C for 90 min, phosphorylated substrate was detected using a phospho-specific antibody, HRP-conjugated secondary antibody, and TMB substrate. The reaction was stopped with an acid stop solution, and absorbance was read at 450 nm on a microplate reader. PKA activity is determined by plotting the reading on the standard curve.

cAMP levels were quantified using the cAMP-Glo™ Max Assay (Promega, TM347). Cells or tissue were treated with agonists as indicated. Samples were prepared according to the manufacturer’s instructions in white, clear-bottom plates. In each well, 40 μl of the tissue lysate was added, followed by 10 μl of the cAMP Detection Solution containing PKA. Following a 20-minute incubation at room temperature, 50 μl of Kinase-Glo® Reagent was added to terminate the PKA reaction and detect residual ATP via luciferase-catalyzed luminescence. Luminescence was measured using a SpectraMax M5 plate reader (Molecular Devices), and cAMP concentrations were determined from a standard curve prepared in parallel.

cGMP content was measured using the Cyclic GMP Complete ELISA Kit (Abcam, ab133052) in the non-acetylated format according to the manufacturer’s instructions. For cell and tissue samples, endogenous phosphodiesterase activity was stopped by treatment with 0.1 M HCl (0.1 g tissue/mL). After 10 min at room temperature, lysates were centrifuged to remove debris, and supernatants were used directly in the assay. Standards, samples, and alkaline phosphatase–conjugated cGMP were added to goat anti-rabbit IgG-coated wells, followed by incubation with cGMP-specific antibody. After washing, pNpp substrate was added, and absorbance was read at 405 nm. cGMP concentrations were calculated from a standard curve prepared in 0.1 M HCl.

### TMT Phosphoproteomic Analysis

Freshly excised left ventricular heart tissues were lysed using Douncing in ice-cold lysis buffer containing 50 mM HEPES (pH 7.5), 6 M guanidine HCl, 10 mM dithiothreitol, protease inhibitor cocktail (S8830, Sigma-Aldrich), and PhosSTOP (04906837001, Roche). An equal amount of protein was used per sample (400 mg) for phosphoproteomic analysis. The digested peptides were labeled with TMT6plex (lot UF288619, Thermo Fisher Scientific) according to the manufacturer’s instructions. Sample fractions were partitioned, with 5% reserved for proteomic analysis and the remaining material completely dried via speed vacuum for subsequent phosphopeptide enrichment. The phosphopeptides were enriched using an adapted immobilized metal affinity chromatography (IMAC) protocol (75). The enriched phosphopeptides underwent a final desalting step using an Empore C18 (2215, 3M) StageTip prior to mass spectrometry (76). Nano-LC/MS/MS analysis was performed using a Dionex rapid separation liquid chromatographic system equipped with an Eclipse instrument (Thermo Fisher Scientific). For data processing, the search criteria included dynamic modification for phosphorylation at serine, threonine, and tyrosine residues. Result validation was performed using the Percolator algorithm against a target-decoy strategy implemented with a concatenated reverse database. Peptide and protein identifications required a false discovery rate (FDR) of <0.01 for high confidence and <0.05 for medium confidence. Abundance values were first corrected for reporter ion isotopic impurity, then normalized to the total summed abundance across all identified peptides within each file’s channel. Data were finalized by exporting protein and phosphopeptide results from Proteome Discoverer and importing them into Perseus (MaxQuant version 1.6.10.43) for subsequent analysis. Group comparisons were performed via Student’s *t*-test (assuming equal variance and requiring two valid total values). The false discovery rate was controlled by calculating the q value using a permutation test. DAVID version 6.8 was employed for enrichment analysis of the differentially phosphorylated proteins. All final data visualization, including heat map generation, was done using R version 3.6.1 (R Foundation for Statistical Computing). Regarding PKA-motif site definition and identification, analyses were limited to class-I localized STY sites. For each site, we extracted a ±7 aa window from the search FASTA and annotated PKA motifs by scanning two basophilic patterns: strict = [RK][RK]xS/T and loose = [RK]..S/T. Only the modified S/T was considered; sites with ambiguous localization or missing windows were excluded. NOS1 KO vs WT (after DOB) was compared on log2 fold-changes with BH-FDR control.

### Immunoprecipitation and Co- Immunoprecipitation

HEK 293 cells expressing WT or mutant PKA regulatory subunits (RIIβ, RIIα), or biotin-switch samples, were solubilized in lysis buffer (in mmol/L, 25 Hepes, 150 NaCl, 5 EDTA, 1.0 % Triton X-100, and protease inhibitors containing 1 PMSF, 2Na3VO4, 10 NaF, 10 µg/mL Aprotinin, 5 Bestatin, 10 µg/mL Leupeptin,2 µg/mL Pepstain A; pH 7.4), followed by centrifugation (12000 rpm, 10 min at 4 °C). The supernatant is then incubated with pre-washed anti-FLAG M2 affinity beads (Sigma-Aldrich) at 4 °C for 2 hours with rotation, and the beads were subsequently washed three times with the lysis buffer. Bound proteins were eluted with 2x sodium dodecyl sulfate loading buffer, followed by centrifugation at maximum speed to pellet the beads, and separated on 8% SDS-PAGE gel. Regarding the dissociation assays, FLAG-RIIβ was immunoprecipitated from HEK293 cells co-expressing PKA catalytic subunit, and eluates were immunoblotted for PKA C. For co-immunoprecipitation of endogenous proteins, ventricular tissue lysates were prepared in the same lysis buffer. We incubated anti-NOS1 Ab overnight at 4 °C with lysates and collected the immune complexes; IgG was included as a negative control.

### Western Blot

Tissues and cells were homogenized in ice-cold lysis buffer containing: 50 mM tris base, 5 mM EDTA, 150 mM NaCl, 1.0% NP-40, 0.1% sodium dodecyl sulfate (SDS), 0.5% sodium deoxycholate 2 mM Na₃VO₄, 1 mM PMSF, 10 mM NaF, aprotinin (10 µg/mL), bestatin (5 mM), leupeptin (10 µg/mL), and pepstatin A (2 µg/mL); PH 7.4. Lysates were incubated on ice for 30 minutes with occasional vortexing and then clarified by centrifugation at 14,000 × g for 10 minutes at 4 °C. Protein concentrations were determined using the BCA assay (Thermo Fisher Scientific, Cat# 23225), and equal amounts of protein were mixed with 2x sodium dodecyl sulfate loading buffer and resolved by SDS-PAGE. Proteins were transferred to PVDF membranes, blocked in 5% non-fat dry milk for 1 h at room temperature, and incubated with primary antibodies overnight at 4 °C. Membranes were washed, incubated with HRP- or IRDye-conjugated secondary antibodies for 1 h at room temperature, developed using ECL, and imaged using a ChemiDoc MP system (Bio-Rad). Band intensities were quantified using Image software and normalized to the appropriate loading control. The following antibodies were used for immunoblotting: anti-PRKAR2B (RIIβ) (ab75993, Abcam); anti-NOS1 (sc-5302, Santa Cruz); anti-NOS3 (sc-376751, Santa Cruz); anti-β1-adrenoceptor (V-19) (sc-568, Santa Cruz); anti-phospho-PKA substrate (RRXS*/T*) (100G7E, #9624, Cell Signaling Technology); anti-SAP97/Dlg1 (610874, BD Transduction Laboratories); anti-phospholamban pSer16 (Abmart, project 20160-1); anti-phospholamban (A010-14, Badrilla); anti-GAPDH (clone 6C5, MAB374, Millipore); anti-PKA RIIα (612242, BD Transduction Laboratories); and anti-PKA C-α (D38C6, #5842, Cell Signaling Technology).

### Immunofluorescence colocalization

Immunofluorescence staining was performed to investigate the subcellular distribution and colocalization of NOS1 with Beta1-adrenoceptors. AVMs were plated on laminin-coated coverslips, fixed with 4% paraformaldehyde for 15 min, and permeabilized with 0.1% Triton X-100 in PBS. After blocking with 5% normal goat serum (NGS) for 1 h, cells were incubated overnight at 4 °C with primary antibodies directed against NOS1 (mouse, 1:100; sc-5302, Santa Cruz Biotechnology) and Beta1-adrenoceptor (rabbit, 1:100; SC-568, Santa Cruz Biotechnology). Following three PBS washes, Alexa Fluor–conjugated secondary antibodies (Invitrogen) were applied for 1 h at room temperature. Cells were mounted, and Images were acquired on a TCS SP8 Falcon confocal microscope (Leica) with a 63× oil-immersion objective under identical laser and gain settings. A sequential scan mode was used with 488-nm excitation for NOS1 (green) and 555-nm excitation for Beta1-adrenoceptors (red). Colocalization analysis was performed using ImageJ with Pearson’s correlation coefficient to quantify overlap between the endogenous NOS1 and Beta1-adrenoceptors, where colocalization was pseudo colored as yellow/orange.

### PKA Sequence and Structure Analysis and Modeling

Crystal structure of PKA RIIβ:C holoenzyme (PDB: 3TNP) was used for structure analysis and *in silico* modeling. We focused on the three cysteine residues: C342, C371 and C386 in the CNB-B domain. C342 and C371 are located in the β-sandwich motif, which cAMP binds, and they are buried into the same hydrophobic pocket. C386 is in the helical subdomain which has conformation changes when comparing the holoenzyme state to the cAMP-bound state of RIIβ. Each cysteine residue was *in silico* modeled with S-Nitrosylation, then followed by the molecular dynamics (MD) simulations. All three cysteine are still in the hydrophobic pocket but S-Nitrosylation enhanced the hydrophobicity of those pockets. All structural figures are made using Pymol (Schrödinger, LLC).

PyMOL was used to create structural representations for the RIIβ:C complex of Protein Kinase A. The complex was obtained from the PDB ID 3TNQ, which is a full-length RIIβ holoenzyme with ADP, magnesium ions, and phosphorylated S122 on the RIIβ subunits (77).

The human genome contains multiple genes that code for proteins containing folds homologous to the cyclic nucleotide binding (CNB) domains found in the regulatory subunits of Protein Kinase A. A total of 40 proteins with collectively 54 CNBs domains were identified using the AlphaFold database, UniProt, and FoldSeek that contain one, two, or three CNB domains, some of which have maintained their functional binding to cAMP and cGMP while others evolved to bind other moieties. Using PyMOL, the CNB domains were separated into individual PDB files and the sequences were converted into FASTA format. To better understand the scope of cysteine nitrosylations in CNB domains, we performed a multiple sequence analysis of every single CNB domain found across the human genome. Initially, Muscle alignment in the Ugene software was used to obtain an approximate alignment. The alignment was subsequently optimized through manual alignment of residues using structural information to ensure that each beta-strand was correctly aligned among all CNB domains. Lastly, the phylogenetic tree was obtained using an IQ-Tree building method using Ugene. For Figure S8, full-length sequences of the regulatory subunits of PKA were aligned (78).

### Human heart RNA-seq Expression Analysis

Processed RNA-seq expression values were obtained from the supplemental gene-level expression matrix provided by Hahn et al (79). For analysis, we generated a mean-centered expression matrix by normalizing each gene to the average expression of that gene across healthy control samples. These normalized values were used to compare transcript levels between control and HFrEF groups.

### Statistical Analysis

All data are expressed as mean ± SEM. Comparisons between two groups were analyzed using an unpaired two-tailed Student’s *t*-test. Paired *t*-tests were applied when comparing baseline and post-stimulation values within the same sample. One-way ANOVA followed by Tukey’s multiple-comparison test was used for comparisons among more than two groups under a single condition, and two-way ANOVA with Sidak’s post hoc test was applied for repeated measures or multiple independent variables. Nonlinear regression was performed to generate concentration–response curves and to compare dose–response relationships. The number of biological replicates (n) for each experiment is indicated in the figure legends. Statistical analysis was conducted using GraphPad Prism (version 10). A value of p < 0.05 was considered statistically significant.

## Abbreviations

AVM: Adult ventricular myocyte
β₁AR β1: adrenergic receptor
β₂AR β2: adrenergic receptor
β3AR β3: adrenergic receptor
BLVRB: biliverdin reductase B
cAMP: Cyclic adenosine monophosphate
cGMP: Cyclic guanosine monophosphate
CL: CL316,243
CNB-B: Cyclic nucleotide-binding domain B
DOB: Dobutamine
EF: Ejection fraction
FRET: Förster resonance energy transfer
GIP: Glucose-dependent insulinotropic polypeptide
GLP1: Glucagon-like peptide 1
GPCR: G protein-coupled receptor
GRK: G protein receptor kinase
INSR: Insulin receptor
ISO: Isoproterenol
KO: Knockout
MI: Myocardial infarction
NO: Nitric oxide
NOS: Nitric oxide synthase
NOS1: Neuronal nitric oxide synthase
NOS3: Endothelial nitric oxide synthase
PKA: Protein kinase A
PKG: Protein kinase
G PLB: Phospholamban
PDE: Phosphodiesterase
PKG1: Protein kinase G type I
PKG2: Protein kinase G type II
RYR: Ryanodine receptor
SCAN: S-nitroso-coenzyme A-assisted nitrosylase
SKF: SKF81297
NECA: 5’-*N*-ethylcarboxamidoadenosine
SERCA: Sarcoplasmic/endoplasmic reticulum Ca²⁺-ATPase
SNO: S-nitrosylation
SNP: Sodium nitroprusside
WT: Wild type

## Author Contributions

SB and YKX conceived the study and designed the overall research plan. SB, ZZ, BX, JW, NC, CZ, WP, CT, ZZ, QW, and YW performed the experiments, analyzed the data, and contributed to the computational work. JH, RP, JS, and SST provided critical conceptual input and scientific guidance. SB and YKX wrote the manuscript. All authors discussed the results and approved the final version of the manuscript.

## Data and materials availability

All data supporting the findings of this study are included in the manuscript and supplementary materials. Additional materials are available upon request.

## List of Supplementary Materials

Fig S1 to S11

Tables S1 and S6

