## Supplemental figures and tables for "S-nitrosylation of protein kinase A is required for its activation by GPCRs"

**Supplementary Materials**

**figure S1. NOS1 deletion reduces β₁-adrenergic receptor–induced PKA activity in heart and brain.**

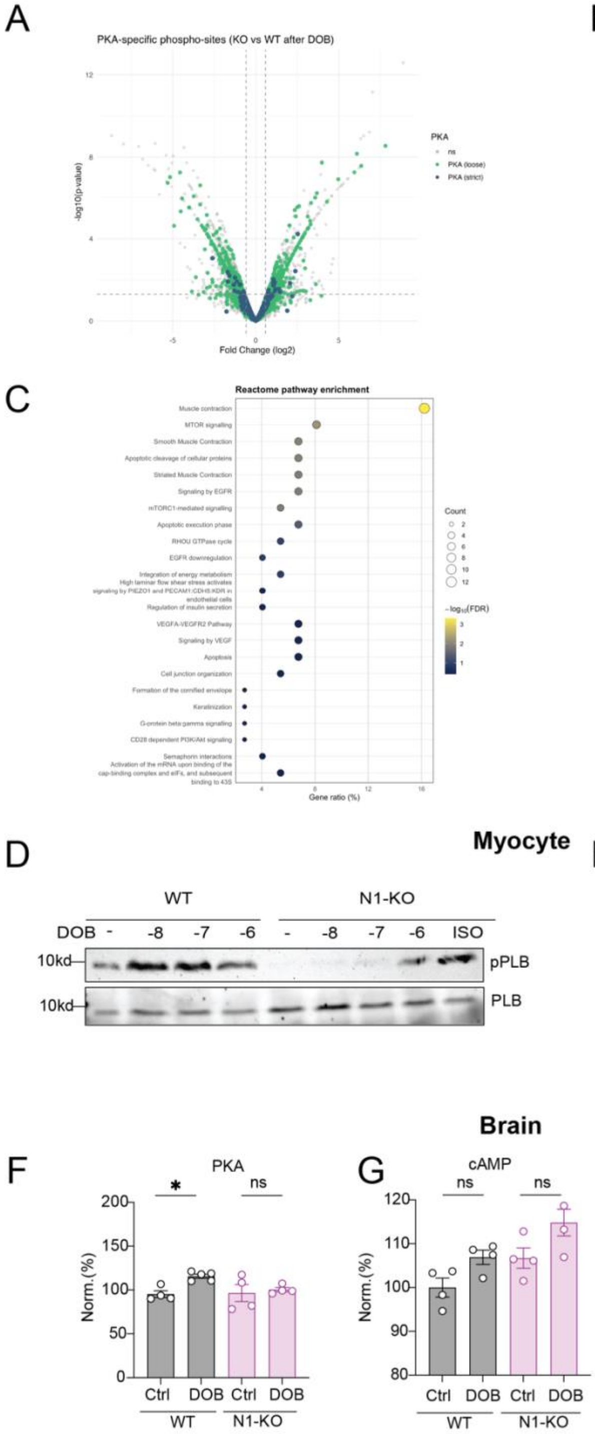


(A) Volcano plot of TMT-based phosphoproteomic analysis of left ventricular tissue from WT and NOS1-KO mice after dobutamine (DOB) stimulation. Each dot represents a quantified phosphosite. The x-axis shows $\log_2$ fold change (KO/WT) and the y-axis shows $-\log_{10}$(p value). Phosphosites containing a stringent PKA consensus motif (R/K-R/K-X-pS/T; "PKA strict") are shown in dark blue, those containing a more relaxed PKA-like motif ("PKA loose") in green, and no significant sites in grey. Vertical dashed lines indicate $\pm\log_2$ (1.5) fold change and the horizontal dashed line marks $p = 0.05$. (B) Bar graph showing the fraction of phosphosites containing a PKA consensus motif (R/K-R/K-X-pS/T) in the total quantified phosphoproteome ("Background") compared with the subset of sites significantly decreased in NOS1-KO versus WT mice after DOB treatment ("KO reduced"). PKA motif sites account for 9.5% of all quantified phosphosites but only 2.7% of KO-reduced sites, indicating that NOS1 deletion preferentially diminishes $\beta_1$-AR–induced phosphorylation at PKA motif sites. (C) Reactome and Gene Ontology enrichment analysis of down-regulated phosphoproteins in DOB-treated NOS1-KO hearts. Bubble size represents gene count; color scale indicates $-\log_{10}$(FDR). Data represent three independent phosphoproteomic replicates per genotype. (D, E) Western blot analysis of pPLB16 in WT and NOS1-KO hearts after DOB stimulation at 10 nM (-8), 100 nM (-7), and 1 μM (-6) (D), with quantification (E). Data are presented as mean ± SEM, n=3, * $P < 0.05$ by One-way ANOVA followed by Tukey test. (F-H) ELISA measurements in brain tissue after DOB stimulation of PKA activity (F), cAMP levels (G) and cGMP levels (H) in NOS1-KO compared with WT. Data are presented as mean ± SEM, n = 4, *$P < 0.05$ by One-way ANOVA followed by Tukey test.

**Figure S2. GPCR–stimulated PKA substrate phosphorylation in heart, brain, and adipose tissue requires NOS.**

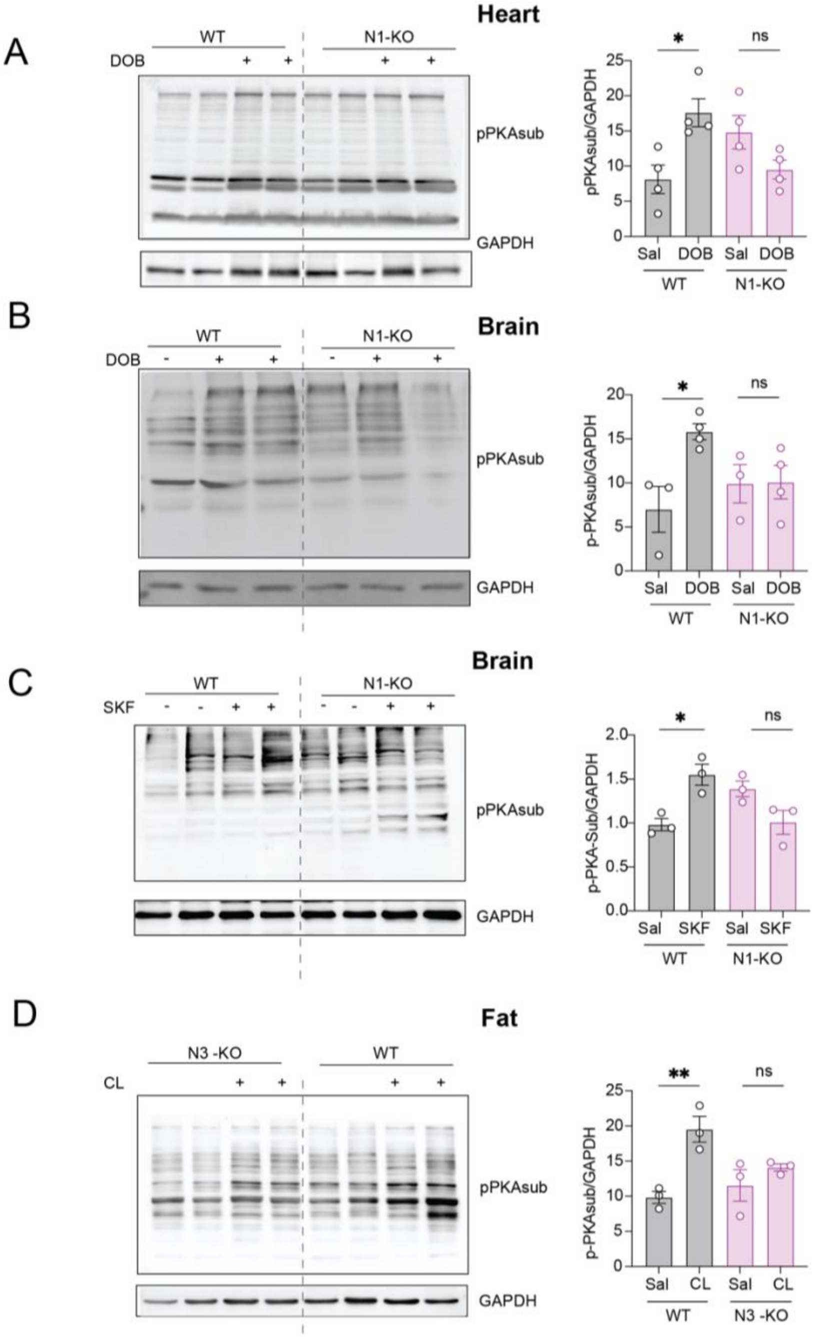


(A–D) Western blot analysis and quantification of ligand-induced PKA substrate phosphorylation across tissues from WT and NOS-KO mice. (A) heart PKA substrate phosphorylation after DOB stimulation in WT and NOS1-KO mice. (B) brain PKA substrate phosphorylation after DOB stimulation in WT and NOS1-KO mice. (C) brain PKA substrate phosphorylation after SKF stimulation in WT and NOS1-KO mice. (D) brown adipose tissue PKA substrate phosphorylation after CL stimulation in WT and NOS1-KO mice. Data are presented as mean ± SEM, n = 3-4. *$P < 0.05$, **$P < 0.01$, by one-way ANOVA with Tukey's multiple comparison test.

**Figure S3. Loss of β₁–NOS1 signaling abolishes β₁-adrenergic contractile responses, while β₂ activation preserves cardiac function.**

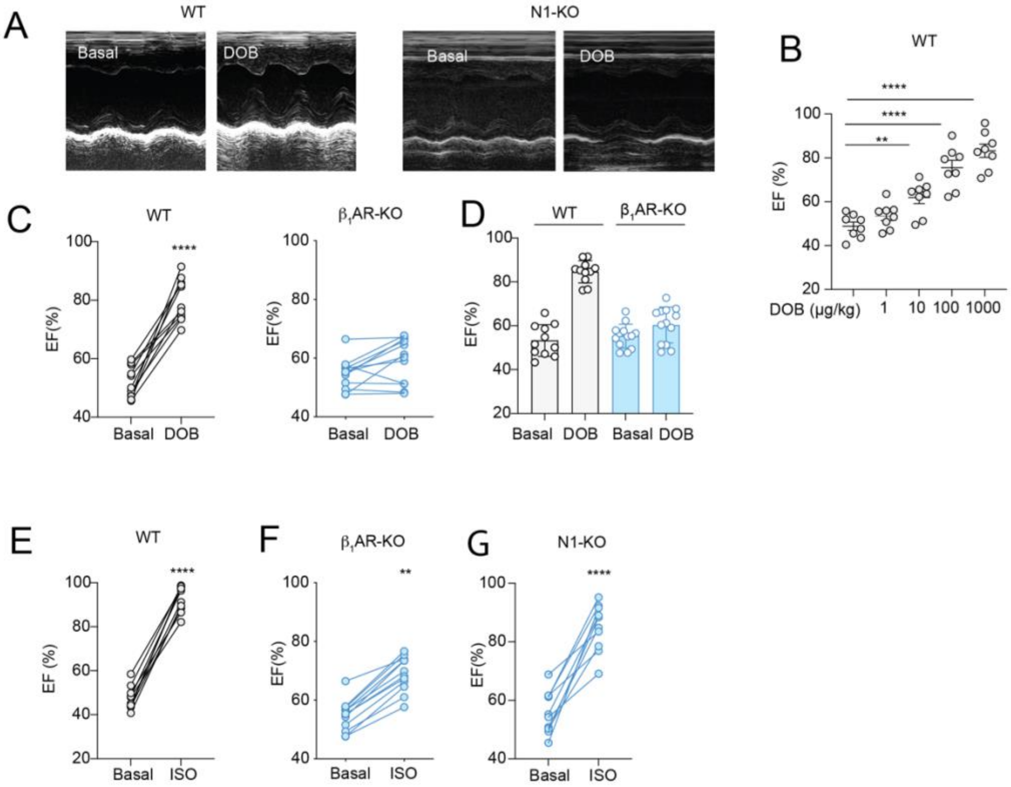


(A, B) Representative echocardiographic images (A) of WT and NOS1-KO mice before and after DOB stimulation, with quantification of ejection fraction (EF%) with varying DOB. ***P* < 0.01, *****P* < 0.0001, by one-way ANOVA with Tukey's multiple comparison test. (C, D) DOB stimulation of ejection fraction in WT and β₁-adrenergic receptor-KO mice. (E-G) ISO stimulation of ejection fraction in WT, β₁AR-KO, and NOS1-KO mice. Data are presented as mean ± SEM from n = 8-10, **P < 0.01, ****P < 0.0001 by Student's paired *t*-test.

**Figure S4. Neither PKG1 nor PKG2 are required for $\beta_1$-adrenergic enhancement of cardiac contractility.**

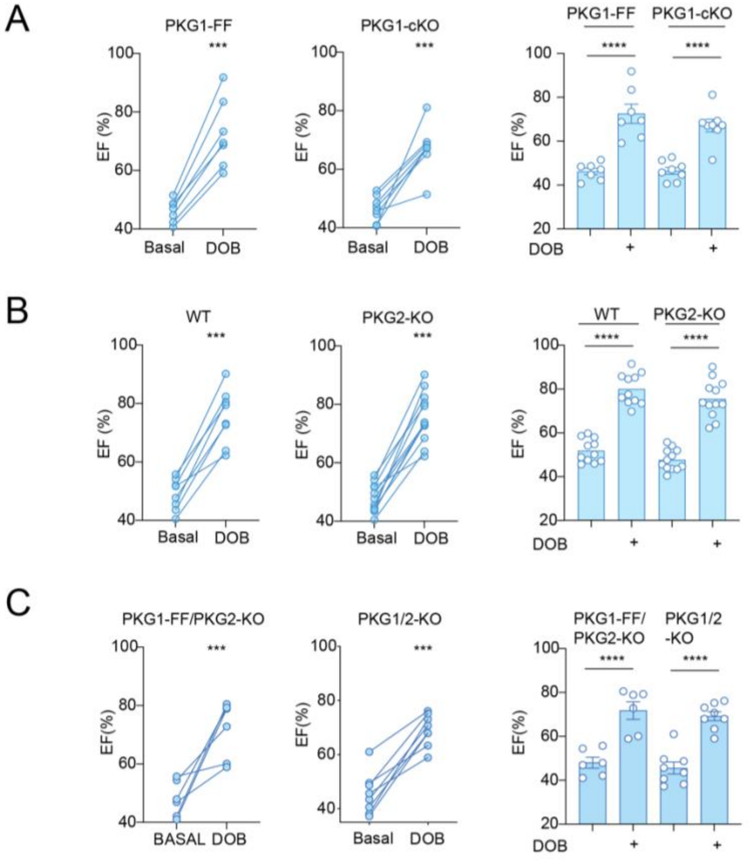


(A) Echocardiography measurement of ejection fraction before and after DOB stimulation in PKG1-cKO mice. (B) Echocardiography measurement of ejection fraction before and after DOB stimulation in PKG2-KO mice. (C) Echocardiography measurement of ejection fraction before and after DOB stimulation in PKG1/2-KO mice. Data are presented as mean ± SEM; n = 7 -10. ***P < 0.001, ****P < 0.0001by Student's paired *t*-test and One-way ANOVA followed by Tukey test.

**Figure S5. Structural proximity and interaction between $\beta_1$-adrenergic receptors and NOS1 in cardiomyocytes.**

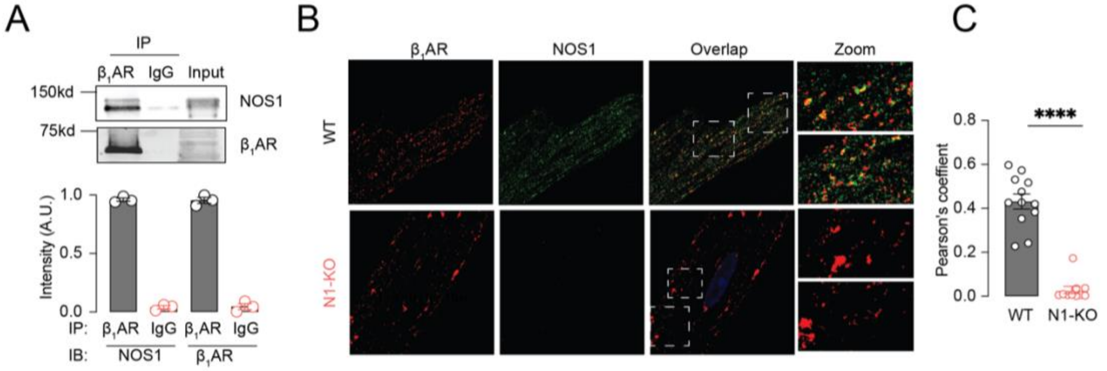


(A) Co-IP of $\beta_1$-adrenergic receptors and NOS1 in WT hearts, ****$P < 0.0001$ by unpaired t test, n = 3. (B) Representative immunofluorescence confocal images of WT and NOS1-KO AVMs stained for $\beta_1$AR (red) and NOS1 (green). WT hearts show extensive co-localization (yellow overlap, right panels), whereas NOS1-KO samples lack the NOS1 signal and overlap. (C) Quantification of $\beta_1$AR–NOS1 immunofluorescence co-localization expressed as Pearson's coefficient demonstrating loss in NOS1-KO AVMs. Data are presented as mean ± SEM; ****$P < 0.0001$ by unpaired *t*-test.

**Figure S6. β-adrenergic stimulation induction of PKA regulatory subunit S-nitrosylation in heart and brain requires NOS1.**

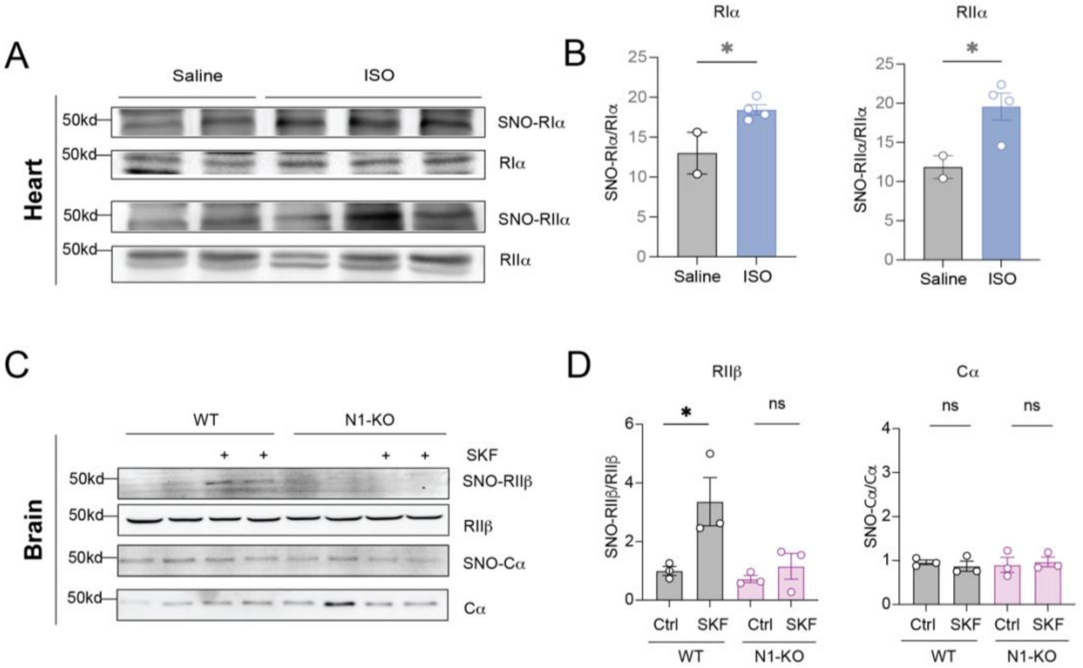


(A–D) Biotin-switch assay-western blot detection of PKA regulatory subunit S-nitrosylation after β-adrenergic stimulation. (A, B) S-nitrosylation of RIα and RIIα subunits in WT hearts from mice perfused with saline or ISO (A), and quantification (B). *P < 0.05 by unpaired Student's *t*-test. (C, D) S-nitrosylation of RIIβ subunits and PKA catalytic subunit-α (Cα) in WT mouse brain treated with saline or SKF (C), and quantification (D). Data are presented as mean ± SEM, n = 3-6, *P < 0.01 by 1-way ANOVA followed by Tukey test.

**Figure S7. NOS1 deletion abolishes $\beta_1$- but not $\beta_2$-adrenergic receptor mediated cGMP signaling.**

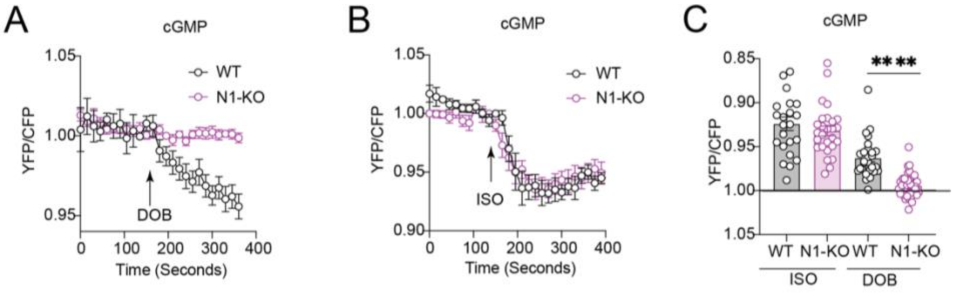


(A-C) Detection of cGMP with FRET biosensor (Gi500) in isolated AVMs from WT and NOS1-KO mice. (A) Representative FRET traces of DOB stimulated cGMP in WT and NOS1-KO cells. (B) Representative FRET traces of ISO stimulated cGMP in WT and NOS1-KO cells. (C) Quantification of peak FRET ratio changes in DOB- and ISO-induced cGMP WT versus NOS1-KO AVMs. Data are presented as mean ± SEM, ****$P < 0.0001$ by One-way ANOVA followed by Tukey test.

**Figure S8. Cysteine residue conservation in PKA regulatory subunits: Sequence alignment and structural modeling.**

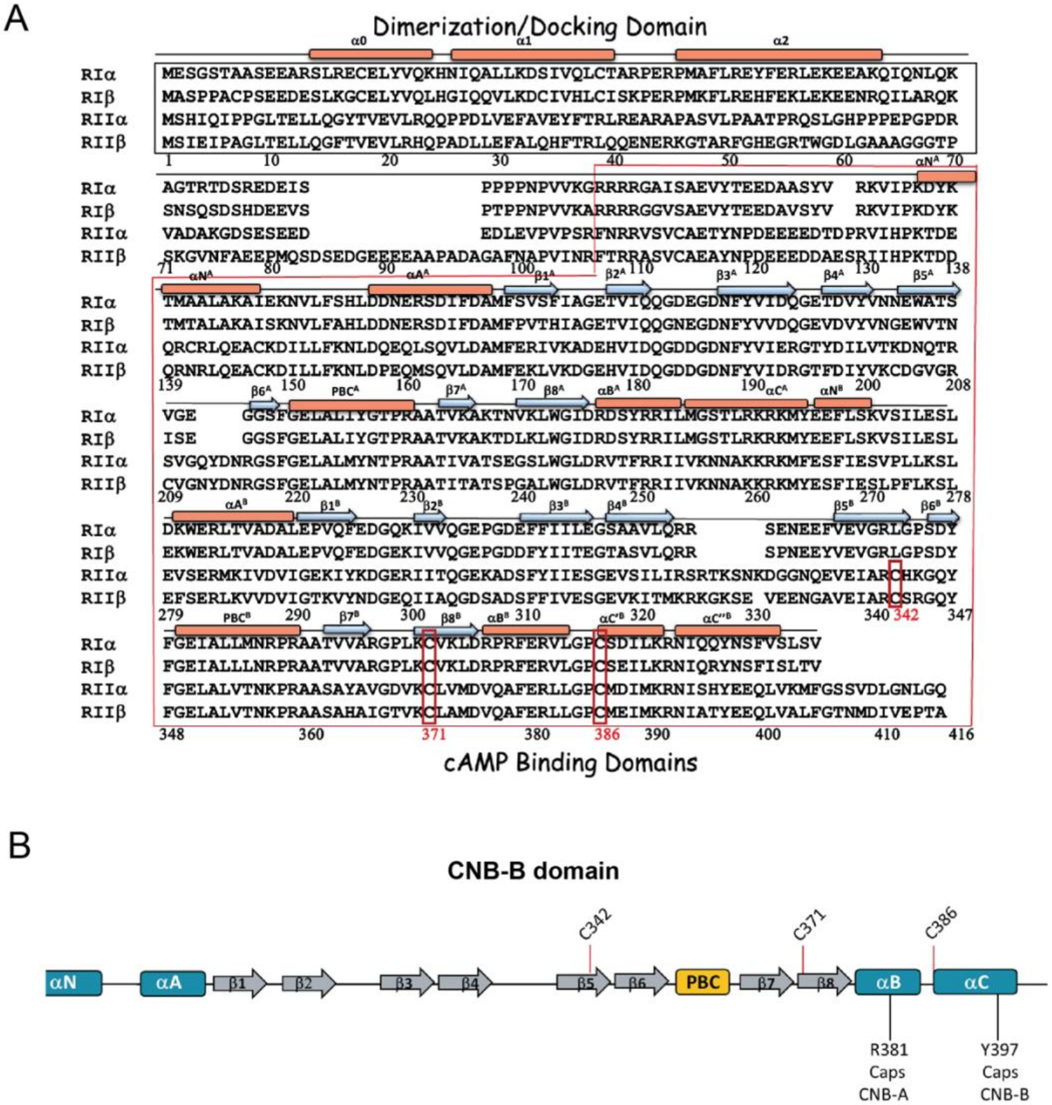


(A) Sequence alignment of PKA regulatory subunits (RIα, RIβ, RIIα, and RIIβ) illustrating conserved structural motifs and domains: the amino terminal Dimerization/Docking (D/D) domain (top black box) mediating subunit dimerization and anchoring interactions, and the carboxyl terminal two cAMP binding domains (bottom black box). Conserved cysteine residues (C342, C371, C386) identified as S-nitrosylated are boxed in red. **(B)** Structural modeling of the second cAMP-binding domain illustrating the positions of Cys371 and Cys386 (red) within hydrophobic pockets surrounded by aromatic and aliphatic residues, suggesting accessibility for S-nitrosylation.

**Figure S9. Sequence analysis and phylogenetic relationships among human CNB-related folds.**

A

| Gene | UniProtID | CNB # | C342 | C371 | C386 |
|---|---|---|---|---|---|
| CNGA1 | P29973 | 1 | L | L | Y |
| CNGA2 | Q16280 | 1 | L | L | Y |
| CNGA3 | Q16281 | 1 | L | L | Y |
| CNGA4 | Q8IV77 | 1 | L | L | Y |
| CNGB1 | Q14028 | 1 | L | L | Y |
| CNGB3 | Q9NQW8 | 1 | L | L | Y |
| CNNM1 | Q9NRU3 | 1 | N | F | C |
| CNNM2 | Q9H8M5 | 1 | A | F | S |
| CNNM3 | Q8NE01 | 1 | N | L | T |
| CNNM4 | Q6P4Q7 | 1 | T | Y | S |
| HCN1 | O60741 | 1 | L | L | Y |
| HCN2 | Q9UL51 | 1 | L | L | Y |
| HCN3 | Q9P1Z3 | 1 | L | L | F |
| HCN4 | Q9Y3Q4 | 1 | L | L | R |
| KCNH1 | O95259 | 1 | L | L | Y |
| KCNH2 | Q12809 | 1 | L | L | Y |
| KCNH3 | Q9ULD8 | 1 | L | L | Y |
| KCNH4 | Q9UQ05 | 1 | L | L | Y |
| KCNH5 | Q8NCM2 | 1 | L | L | Y |
| KCNH6 | Q9H252 | 1 | L | L | Y |
| KCNH7 | Q9NS40 | 1 | L | L | Y |
| KCNH8 | Q96L42 | 1 | L | L | Y |
| KDM2B | F5H7T7 | 1 | L | V | |
| KGP1 | Q13976 | 1 | M | L | T |
| KGP1 | Q13976 | 2 | L | C | L |
| KGP2 | Q13237 | 1 | I | T | T |
| KGP2 | Q13237 | 2 | L | C | F |
| PLPL6 | Q8IY17 | 1 | V | V | Y |
| PLPL6 | Q8IY17 | 2 | A | F | Q |
| PLPL6 | Q8IY17 | 3 | Y | L | Y |
| PLPL7 | Q6ZV29 | 1 | V | I | Y |
| PLPL7 | Q6ZV29 | 2 | T | F | Q |
| PLPL7 | Q6ZV29 | 3 | Y | L | Y |
| POPD1 | Q8NE79 | 1 | I | F | E |
| POPD2 | Q9HBU9 | 1 | I | Y | E |
| POPD3 | Q9HBV1 | 1 | I | Y | H |
| PRKAR1A | P10644 | 1 | V | L | S |
| PRKAR1A | P10644 | 2 | L | C | C |
| PRKAR1B | P31321 | 1 | I | L | S |
| PRKAR1B | P31321 | 2 | L | C | C |
| PRKAR2A | P13861 | 1 | Y | L | N |
| PRKAR2A | P13861 | 2 | C | C | C |
| PRKAR2B | P31323 | 1 | Y | L | N |
| PRKAR2B | P31323 | 2 | C | C | C |
| RAPGEF2 | Q9Y4G8 | 1 | L | F | V |
| RAPGEF3 | O95398 | 1 | L | F | V |
| RAPGEF4 | Q8WZA2 | 1 | L | F | V |
| RAPGEF4 | Q8WZA2 | 2 | L | L | Y |
| RAPGEF6 | Q8TEU7 | 1 | L | F | V |
| RAPGEF6 | Q8TEU7 | 2 | V | M | Q |
| SL9C1 | Q4G0N8 | 1 | M | T | C |
| SL9C1 | Q4G0N8 | 2 | Y | V | N |
| SL9C2 | Q5TAH2 | 1 | C | A | F |
| SL9C2 | Q5TAH2 | 2 | Y | L | T |
| Consensus Residue | | | L | L | Y |
| Conservation % | | | 51.85% | 51.85% | 39.62% |

B

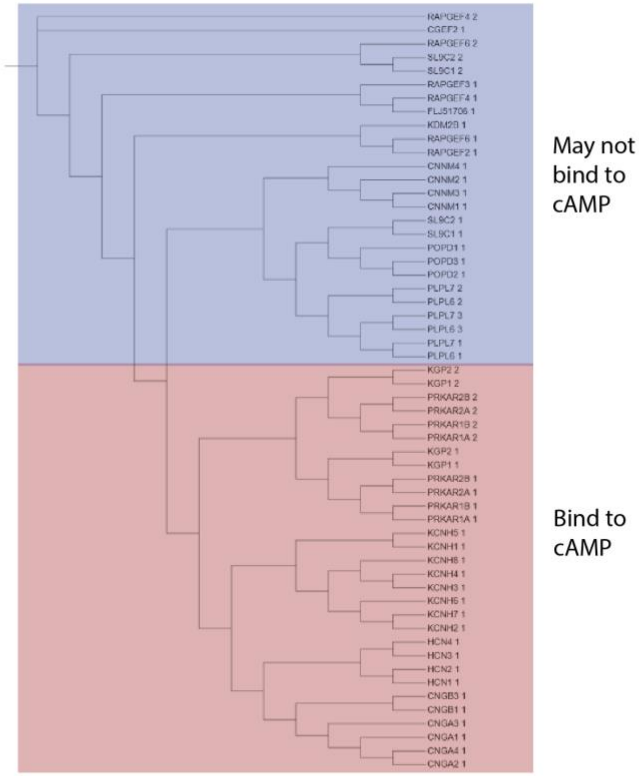


(A) Cysteine triad conservation across human CNB-related folds (using RIIβ numbering). In PKA, cysteine residues are conserved only within the second CNB domains. (B) phylogenetic sequence alignment of representative human CNB-related protein domains (UniProt ID followed by CNB number).

**Figure S10. S-Nitrosylation enhances the hydrophobicity of cysteine residue in cAMP binding pockets of RII subunit and activation of PKA.**

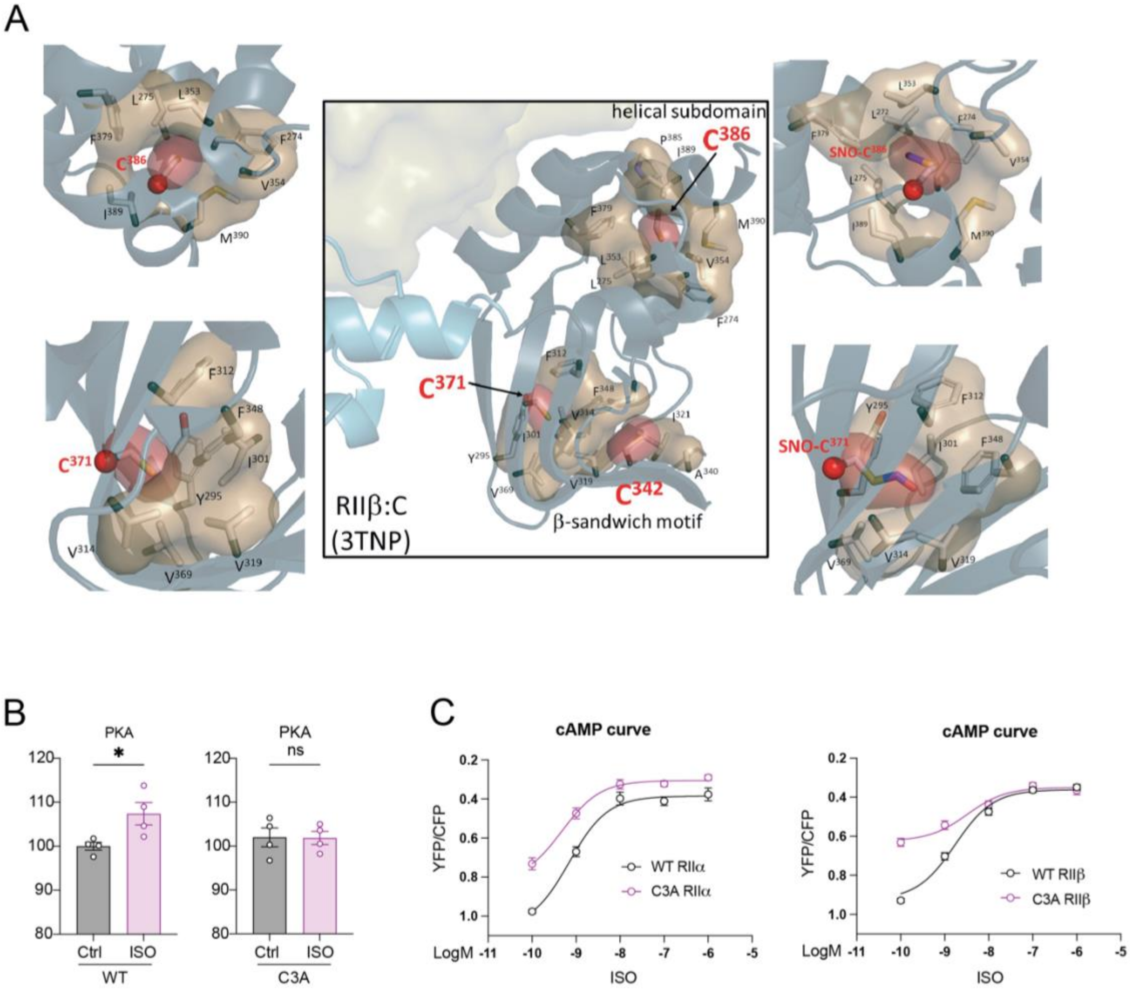


(A) All three cysteine residues are located in hydrophobic pockets in the CNB-B domain of the PKA RIIβ:C holoenzyme structure (Middle panel). Each CNB domain contains two subdomains. Two cysteines (C342 and C371) are part of the β-sandwich motif where cAMP docks, while C386 is in the helical subdomain. The phosphate-binding cassette (PBC) is part of the β-sandwich motif while the two hydrophobic residues that cap cAMP bound to CNB-A (R381) and CNB-B (Y397) are located in the C-terminal B/C helices of the CNB-B domain. Left panels show the zoom-in view of C371 and C386. The sand-colored shells show the hydrophobic residues that surrounding each cysteine (red shell). The *in silico* modeling of C371 and C386 with S-Nitrosylation, shown on the right panels, indicated that there is no steric clash. The S-Nitrosylation simply enhances the hydrophobicity of each pocket. There is also no steric clash with cAMP binding (data not shown). (B) Co-immunoprecipitation of overexpressed PKA catalytic subunit with WT or triple-mutant RIIβ in HEK293 cells treated with ISO. PKA activity was assessed by ELISA in the immunoprecipitated complexes. Data are presented as mean ± SEM, n = 4. *P < 0.05 by unpaired Student's t test. (C) FRET H187 biosensor analysis of cAMP dose–response curves after ISO stimulation in HEK293 cells overexpressing WT or triple-mutant (C342A/C371A/C386A) RIIβ or equivalent RIIα mutants, demonstrating comparable (or slightly elevated) cAMP responses in the cells expressing mutant RII subunits. The log $EC_{50}$ values for cAMP were −9.22 and −9.43 for WT and RIIα triple mutant, and −8.76 and −8.48 for WT and RIIβ triple mutant, respectively. Dose-response curves were analyzed by nonlinear regression. Data are mean ± SEM, n = 4-5.

**Figure S11. SCAN is necessary for GPCR-mediated activation of PKA.**

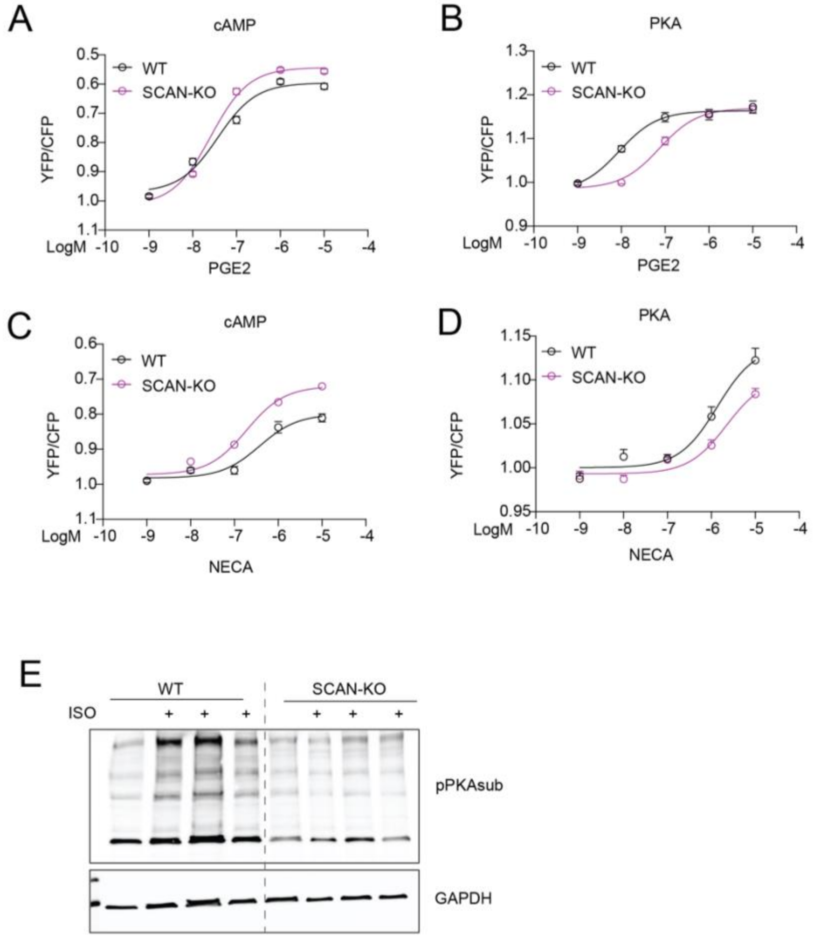


(A, B) FRET-based cAMP (panel A, H187 sensor) and PKA activity (panel B, AKAR3 sensor) dose-response analysis in WT and SCAN-KO HEK293T cells stimulated by $PGE_2$. (C, D) FRET-based cAMP (panel C, H187 sensor, log $EC_{50} = -6.43$ for WT and $-6.73$ for SCAN-KO) and PKA activity (panel D, AKAR3 sensor, log $EC_{50} = -5.86$ for WT and $-5.65$ for SCAN-KO) dose-response analysis in WT and SCAN-KO HEK293T cells stimulated by NECA. Dose-response curves generated by nonlinear regression analysis. (E) Western blot of PKA substrate phosphorylation after ISO (1 nM) stimulation in normal and SCAN-KO HEK293 cells. Quantification shown in Fig. 4F for n = 4-5.

**Figure S12. Altered gene expression in human HFrEF patients.**

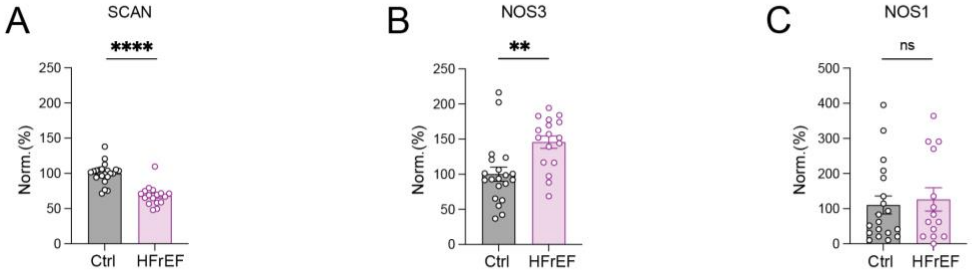


(A–C) Human cardiac RNA sequencing data from control and HFrEF samples showing SCAN (A), NOS3 (B), and NOS1 (C) transcript levels. Data are presented as mean ± SEM; n = 17. **P < 0.01, ****P < 0.0001 by unpaired Student's t-test.

**table S1. Dobutamine-induced cardiac functional enhancement in WT but blunted in β1AR KO mice.**

Cardiac function before and after Dobutamine treatment was assessed by 2D M-mode short-axis echocardiography. WT and β1AR KO mice were injected with dobutamine (DOB; i.p., 400 ng/g, N = 11 per group). Data are expressed as mean ± SEM. Paired t-test was used to compare Baseline and Post-DOB values within each genotype. *$p < 0.05$, **$p < 0.01$, and ***$p < 0.001$ versus Baseline (WT). DOB significantly enhanced systolic function in WT mice, as indicated by increased EF and FS, whereas this inotropic response was attenuated in β1AR KO mice.

| Group | WT | | β1AR KO | |
|---|---|---|---|---|
| | Baseline | DOB | Baseline | DOB |
| HR (bpm) | 496.2 ± 9.75 | 492.4 ± 4.23 | 338.7 ± 8.88 | 322.2 ± 10.31 |
| IVS; d(mm) | 0.69 ± 0.04 | 0.83 ± 0.05** | 0.71 ± 0.02 | 0.75 ± 0.04 |
| IVS; s(mm) | 1.04 ± 0.03 | 1.30 ± 0.04** | 1.06 ± 0.03 | 1.16 ± 0.03 |
| LVID; d(mm) | 3.86 ± 0.04 | 3.52 ± 0.07** | 3.55 ± 0.07 | 3.53 ± 0.07 |
| LVID; s(mm) | 2.88 ± 0.08 | 2.04 ± 0.007*** | 2.6 ± 0.07 | 2.5 ± 0.07 |
| LVPW; d(mm) | 0.71 ± 0.02 | 0.75 ± 0.02 | 0.83 ± 0.03 | 0.83 ± 0.04 |
| LVPW; s(mm) | 0.97 ± 0.04 | 1.28 ± 0.04*** | 1.04 ± 0.04 | 1.09 ± 0.04 |
| EF % | 50.87 ± 2.29 | 73.94 ± 1.25*** | 54.51 ± 1.64 | 58.69 ± 2.31 |
| FS % | 25.53 ± 1.41 | 42.0 ± 1.06*** | 27.71 ± 1.07 | 30.31 ± 1.50 |
| LV Mass (mg) | 92.71 ± 5.04 | 92.95 ± 3.63 | 81.13 ± 5.54 | 83.63 ± 5.82 |
| LV Vol; d(uL) | 64.26 ± 1.57 | 51.60 ± 2.40** | 52.97 ± 2.60 | 52.09 ± 2.32 |
| LV Vol; s(uL) | 31.69 ± 2.10 | 13.52 ± 1.10*** | 23.95 ± 1.66 | 20.97 ± 1.72 |

**table S2. β1AR stimulation enhances cardiac contraction in WT, NOS3KO, but not NOS1KO mice.**

Cardiac function before and after dobutamine (DOB) treatment was measured by 2D M-mode short-axis echocardiography. WT, NOS1KO, and NOS3KO mice were treated with DOB (i.p., 400 ng/g, N = 11). Data shown are mean values ± SEM. Paired t-tests were performed to compare Baseline and DOB within each genotype. *p<0.05, **p<0.01, ***p<0.001 when compared with Baseline (WT); #p<0.05, ##p<0.01, ####p<0.001 when compared with Baseline (NOS3KO).

| Group | WT | | NOS1KO | | NOS3KO | |
|---|---|---|---|---|---|---|
| | Basal | DOB | Basal | DOB | Basal | DOB |
| HR (bpm) | 470.6 ± 6.047 | 516.4 ± 9.325 | 440.0 ± 15.87 | 414.1 ± 14.09 | 470.2 ± 6.928 | 481.9 ± 10.14 |
| IVS; d (mm) | 0.6188 ± 0.01465 | 0.7270 ± 0.02706 | 0.6161 ± 0.01795 | 0.6460 ± 0.01949 | 0.7389 ± 0.08866 | 0.8629 ± 0.07152 |
| IVS; s (mm) | 0.9119 ± 0.04322 | 1.274 ± 0.06134* | 0.8632 ± 0.02695 | 0.9506 ± 0.02018 | 1.077 ± 0.04116 | 0.9506 ± 0.03546 |
| LVID; d (mm) | 3.315 ± 0.1747 | 2.915 ± 0.1618 | 3.452 ± 0.07278 | 3.342 ± 0.08977 | 3.611 ± 0.07392 | 3.134 ± 0.07578 |
| LVID; s (mm) | 2.481 ± 0.2323 | 1.366 ± 0.1727*** | 2.594 ± 0.08450 | 2.354 ± 0.1346 | 2.582 ± 0.09015 | 1.435 ± 0.08236### |
| LVPW; d (mm) | 0.6077 ± 0.03244 | 0.7438 ± 0.03614 | 0.6138 ± 0.02821 | 0.6086 ± 0.02280 | 0.7784 ± 0.04445 | 0.9182 ± 0.04946 |
| LVPW; s (mm) | 0.9005 ± 0.05027 | 1.278 ± 0.08385* | 0.8833 ± 0.04781 | 0.9115 ± 0.05684 | 1.142 ± 0.03742 | 1.550 ± 0.04076# |
| EF % | 51.68 ± 5.345 | 84.99 ± 3.061**** | 50.35 ± 1.993 | 57.57 ± 3.744 | 57.74 ± 2.503 | 84.96 ± 1.643#### |
| FS % | 26.33 ± 3.365 | 54.24 ± 3.738**** | 25.02 ± 1.174 | 30.07 ± 2.591 | 27.99± 1.579 | 53.40 ± 1.995#### |
| LV Mass (mg) | 60.85 ± 5.197 | 62.99 ± 4.473 | 64.75 ± 2.50 | 56.04 ± 2.912 | 92.10 ± 4.074 | 92.87 ± 5.122 |
| LV Vol; d (μL) | 46.00 ± 5.685 | 33.79 ± 4.385 | 22.95 ± 2.366 | 43.48 ± 5.904 | 57.08 ± 2.753 | 40.90 ± 2.230#### |
| LV Vol; s (μL) | 24.21 ± 5.167 | 5.822 ± 1.752*** | 24.90 ± 1.971 | 20.30 ± 8.458 | 26.08 ± 2.276 | 6.300 ± 0.8654#### |

**table S3. β1AR stimulation enhanced cardiac contraction in PKG1-F/F and PKG1-cKO mice.**

Cardiac function before and after drug treatment was measured by 2D M-mode short-axis echocardiography. PKG1-F/F and PKG1-KO mice were treated with Dobutamine (DOB *i.p.,* 400 ng/g, N = 8 per group). Data shown are mean values ± SEM. Paired *t*-test was performed to analyze the differences Basal and After conditions. * p<0.05, **p<0.01, and ***p<0.001 when compared with PKG1-F/F Baseline (DOB), and $^{@}$p<0.05 $^{@@}$p<0.01, $^{@@@}$p<0.001, and $^{@@@@}$p<0.001 when compared with PKG1-cKO Baseline (DOB).

| Group | PKG1-F/F | | PKG1-cKO | |
|---|---|---|---|---|
| | Baseline | DOB | Baseline | DOB |
| HR (bpm) | 442.3 ± 6.42 | 539.7 ± 7.58*** | 461 ± 12.69 | 541.4 ± 14.79$^{@}$ |
| IVS; d(mm) | 0.73 ± 0.07 | 0.80 ± 0.05*** | 0.66 ± 0.04 | 0.74 ± 0.05$^{@@@}$ |
| IVS; s(mm) | 0.81 ± 0.04 | 1.09 ± 0.04*** | 0.81 ± 0.03 | 1.01 ± 0.05$^{@@@}$ |
| LVID; d(mm) | 3.65 ± 0.07 | 3.14 ± 0.06*** | 3.80 ± 0.10 | 3.50 ± 0.15$^{@@@}$ |
| LVID; s(mm) | 2.90 ± 0.07 | 1.83 ± 0.14*** | 3.07 ± 0.09 | 2.23 ± 0.15$^{@@@}$ |
| LVPW; d(mm) | 0.73 ± 0.04 | 0.71 ± 0.06*** | 0.75 ± 0.03 | 0.80 ± 0.03$^{@@@}$ |
| LVPW; s(mm) | 0.94 ± 0.05 | 1.22 ± 0.07*** | 0.95 ± 0.03 | 1.30 ± 0.07$^{@@@}$ |
| EF % | 42.64 ± 1.25 | 72.50 ± 3.34**** | 40.44 ± 1.06 | 67.17 ± 2.84$^{\$\$\$\$}$ |
| FS % | 20.51 ± 0.70 | 41.71 ± 4.30**** | 19.35 ± 0.58 | 36.74 ± 2.12$^{@@@}$ |
| LV Mass (mg) | 89.53 ± 5.67 | 74.03 ± 6.30*** | 91.41 ± 6.16 | 91.74 ± 7.93$^{@@@}$ |
| LV Vol; d(μL) | 56.49 ± 2.79 | 39.30 ± 1.80*** | 62.80 ± 3.91 | 52.05 ± 5.10$^{@@@}$ |
| LV Vol; s(μL) | 32.43 ± 1.84 | 10.79 ± 1.85*** | 37.41 ± 2.38 | 17.67 ± 3.06$^{@@@}$ |

**table S4. β1AR stimulation enhanced cardiac contraction in WT and PKG2 KO mice.** Cardiac function before and after drug treatment was measured by 2D M-mode short-axis echocardiography. WT and PKG2 KO mice were treated with Dobutamine (DOB *i.p.,* 400 ng/g, N = 8 for WT and N = 11 for PKG2 KO). Data shown are mean values ± SEM. Paired *t*-test was performed to analyze the differences Basal and After conditions. * p<0.05, **p<0.01, and ***p<0.001 when compared with WT Baseline (DOB), and $^{\$\$}$p<0.01, $^{\$\$\$}$p<0.001, and $^{\$\$\$}$p<0.001 when compared with PKG2 KO Baseline (DOB).

| Group | WT | | PKG2-KO | |
|---|---|---|---|---|
| | Baseline | DOB | Baseline | Stimulation |
| HR (bpm) | 467.6 ± 6.42 | 502.9 ± 7.97* | 421.3 ± 20.59 | 466.4 ± 22.80$^{\$}$ |
| IVS; d(mm) | 0.62 ± 0.05 | 0.72 ± 0.04* | 0.57 ± 0.03 | 0.67 ± 0.04$^{\$\$}$ |
| IVS; s(mm) | 0.70 ± 0.05 | 0.91 ± 0.04** | 0.80 ± 0.04 | 1.05 ± 0.04$^{\$\$\$}$ |
| LVID; d(mm) | 3.19 ± 0.16 | 2.88 ± 0.15**** | 3.72 ± 0.09 | 3.16 ± 0.11$^{\$\$\$}$ |
| LVID; s(mm) | 2.43 ± 0.14 | 1.62 ± 0.12**** | 2.87 ± 0.08 | 1.87 ± 0.12$^{\$\$\$\$}$ |
| LVPW; d(mm) | 0.76 ± 0.03 | 0.79 ± 0.03 | 0.63 ± 0.06 | 0.71 ± 0.03 |
| LVPW; s(mm) | 0.99 ± 0.05 | 1.36 ± 0.06*** | 0.85 ± 0.07 | 1.18 ± 0.07$^{\$\$\$\$}$ |
| EF % | 48.86 ± 1.91 | 75.55 ± 3.34**** | 46.57 ± 1.49 | 73.12 ± 2.51$^{\$\$\$\$}$ |
| FS % | 23.9 ± 1.12 | 43.67 ± 3.16*** | 22.85 ± 0.89 | 41.30 ± 2.07$^{\$\$\$\$}$ |
| LV Mass (mg) | 66.65 ± 4.03 | 64.42 ± 4.19 | 73.20 ± 9.0 | 66.31 ± 4.99 |
| LV Vol; d(μL) | 41.92 ± 5.38 | 32.62 ± 4.19*** | 59.40 ± 3.63 | 40.59 ± 3.40$^{\$\$\$}$ |
| LV Vol; s(μL) | 21.62 ± 3.19 | 8.0 ± 1.44*** | 31.81 ± 2.34 | 11.53 ± 1.93$^{\$\$\$\$}$ |

**table S5. β1AR stimulation increases cardiac contraction in WT mice, but not in NOS1KO mice, where pretreatment with SNP restores the DOB effect.**

Cardiac function before and after dobutamine (DOB) treatment was measured by 2D M-mode short-axis echocardiography. WT and NOS1KO mice were treated with DOB (i.p., 400 ng/g, N = 8), SNP (N = 8) (100 ng/g), and DOB + SNP (N = 8). Data shown are mean values ± SEM. One-way ANOVA was performed to compare the Baseline and DOB within each genotype. *p<0.05, **p<0.01, ***p<0.001 when compared with Baseline (WT); #p<0.05, ##p<0.01, ####p<0.001 when compared with Baseline (NOS1KO).

| Group | WT | | NOS1KO | | | |
|---|---|---|---|---|---|---|
| | Basal | DOB | Basal | DOB | SNP | SNP+DOB |
| HR (bpm) | 460.3 ± 17.32 | 461.0 ± 17.44 | 442.0 ± 18.94 | 398.6 ± 11.67 | 445.4 ± 10.42 | 357.1 ± 11.08## |
| IVS; d (mm) | 0.70 ± 0.04 | 0.74 ± 0.05 | 0.62 ± 0.02 | 0.65 ± 0.02 | 0.54 ± 0.02 | 0.64 ± 0.02 |
| IVS; s (mm) | 0.98 ± 0.04 | 1.32 ± 0.05**** | 0.87 ± 0.03 | 0.97 ± 0.02 | 0.84 ± 0.01 | 1.06 ± 0.03### |
| LVID; d (mm) | 3.84 ± 0.09 | 3.54 ± 1.0 | 3.45 ± 0.09 | 3.30 ± 0.11 | 3.74 ± 0.06 | 3.45 ± 0.07 |
| LVID; s (mm) | 2.81 ± 0.08 | 1.72 ± 0.10**** | 2.62 ± 0.10 | 2.32 ± 0.16 | 2.72 ± 0.07 | 1.98 ± 0.06### |
| LVPW; d (mm) | 0.49 ± 0.03 | 0.59 ± 0.05 | 0.60 ± 0.03 | 0.60 ± 0.03 | 0.56 ± 0.02 | 0.64 ± 0.01 |
| LVPW; s (mm) | 0.82 ± 0.06 | 1.35 ± 0.06**** | 0.84 ± 0.04 | 0.89 ± 0.06 | 0.84 ± 0.04 | 1.14 ± 0.02## |
| EF % | 53.14 ± 1.78 | 82.96 ± 2.16**** | 49.68 ± 2.16 | 53.59 ± 3.19 | 54.31 ± 1.07 | 74.80 ± 0.85#### |
| FS % | 26.97 ± 1.11 | 51.47 ± 2.43**** | 24.29 ± 1.30 | 30.41 ± 3.03 | 27.31 ± 0.68 | 42.69 ± 0.71#### |
| LV Mass (mg) | 73.97 ± 2.97 | 74.73 ± 4.27 | 64.01 ± 2.97 | 56.08 ± 3.58 | 64.39 ± 2.47 | 68.15 ± 3.10 |
| LV Vol; d (μL) | 63.95 ± 3.50 | 52.80 ± 3.51 | 49.64 ± 3.01 | 44.81 ± 3.34 | 59.71 ± 2.37 | 49.44 ± 2.30 |
| LV Vol; s (μL) | 30.08 ± 2.27 | 9.05 ± 1.35**** | 25.60 ± 2.36 | 19.73 ± 3.03 | 27.66 ± 1.60 | 12.54 ± 0.88### |

**table S6. Cardiac function of WT MI mice before and after dobutamine, SNP or DOB+SNP treatment.**

MI mice were subjected to echocardiography measurement before and after dobutamine (N = 10) (400 ng/g), SNP (N = 10) (100 ng/g), DOB + SNP (N = 10) intraperitoneal injection. Cardiac function assessed by 2D M-mode. Data shown are mean values ± SEM, * $p<0.05$ when compared with Baseline (DOB), $^{\$}p<0.05$ when compared with Baseline (DOB+SNP). $^{\#}p<0.05$ when compared with Baseline in Shan (Baseline in MI). One-way ANOVA was performed to analyze changes between different groups.

| Group | Sham | | | | MI | | | |
|---|---|---|---|---|---|---|---|---|
| | Baseline | Dob | SNP | Dob+SNP | Baseline | Dob | SNP | Dob+SNP |
| EF (%) | 48.44 ± 1.922 | 64.96 ± 2.951* | 49.34 ± 4.048 | 63.80 ± 3.274$^{\$}$ | 33.30 ± 3.282$^{\#}$ | 38.26 ± 3.684 | 36.95 ± 2.313 | 57.26 ± 3.642$^{\$}$ |
| FS (%) | 23.14 ± 0.7450 | 35.16 ± 2.192* | 22.88 ± 1.718 | 34.63 ± 2.474$^{\$}$ | 12.10 ± 1.129$^{\#}$ | 16.87 ± 1.875 | 17.74 ± 1.210 | 28.66 ± 2.211$^{\$}$ |
| Heart Rate (BPM) | 477.8 ± 24.12 | 473.6 ± 15.12 | 410.1 ± 16.23 | 418.2 ± 26.10 | 481.6 ± 10.12 | 412.8 ± 14.28 | 416.7 ± 20.04 | 427.4 ± 15.35 |
| Diameter;s (mm) | 2.974 ± 0.07786 | 2.404 ± 0.1207 | 2.972 ± 0.1481 | 2.407 ± 0.1423 | 3.786 ± 0.1856$^{\#}$ | 3.628 ± 0.1887 | 3.496 ± 0.2090 | 2.688 ± 0.2092$^{\$}$ |
| Diameter;d (mm) | 3.916 ± 0.06202 | 3.626 ± 0.07362 | 3.956 ± 0.06871 | 3.663 ± 0.1014 | 4.504 ± 0.1424$^{\#}$ | 4.433 ± 0.1410 | 4.355 ± 0.1526 | 3.915 ± 0.1750$^{\$}$ |
| Volume;s (uL) | 34.63 ± 2.167 | 20.98 ± 2.623 | 35.50 ± 3.714 | 21.49 ± 3.107 | 64.15 ± 7.411$^{\#}$ | 58.26 ± 6.900 | 53.57 ± 6.863 | 29.45 ± 4.752$^{\$}$ |
| Volume;d (uL) | 66.81 ± 2.523 | 55.69 ± 2.717 | 68.50 ± 2.759 | 57.36 ± 3.735 | 94.13 ± 6.893$^{\#}$ | 90.86 ± 6.622 | 87.18 ± 6.982 | 68.54 ± 6.576 |
| Stroke Volume (uL) | 32.18 ± 1.304 | 34.71 ± 1.432 | 33.00 ± 1.590 | 35.88 ± 1.786 | 27.08 ± 2.233 | 32.60 ± 2.191 | 33.60 ± 2.511 | 39.09 ± 3.056$^{\$}$ |
| Cardiac Output (mL/min) | 15.96 ± 0.8788 | 16.55 ± 1.069 | 13.65 ± 1.052 | 14.23 ± 1.225 | 13.68 ± 1.263 | 12.87 ± 1.092 | 14.27 ± 1.674 | 16.80 ± 1.527 |
| LV Mass (mg) | 74.55 ± 3.996 | 75.10 ± 3.665 | 78.22 ± 4.057 | 74.95 ± 4.278 | 92.89 ± 6.813 | 92.29 ± 7.817 | 87.51 ± 10.98 | 85.75 ± 8.894 |
| LV Mass Cor (mg) | 59.64 ± 3.197 | 60.08 ± 2.932 | 62.57 ± 3.246 | 59.96 ± 3.422 | 74.32 ± 5.450 | 73.83 ± 6.253 | 70.01 ± 8.784 | 68.60 ± 7.115 |
| LVAW;s (mm) | 0.8145 ± 0.04171 | 1.006 ± 0.05428 | 0.8272 ± 0.05047 | 1.009 ± 0.05059 | 0.6424 ± 0.06871 | 0.7338 ± 0.06487 | 0.7006 ± 0.1094 | 0.9202 ± 0.1091 |
| LVAW;d (mm) | 0.5602 ± 0.03076 | 0.6168 ± 0.02326 | 0.5778 ± 0.03618 | 0.6281 ± 0.03312 | 0.4888 ± 0.04377 | 0.5123 ± 0.03320 | 0.4881 ± 0.06027 | 0.5700 ± 0.05507 |
| LVPW;s (mm) | 0.9039 ± 0.04991 | 1.130 ± 0.05376 | 0.9017 ± 0.07920 | 1.092 ± 0.05213 | 0.8700 ± 0.04928 | 0.8943 ± 0.03645 | 0.9023 ± 0.08113 | 1.153 ± 0.08899$^{\$}$ |
| LVPW;d (mm) | 0.5980 ± 0.02593 | 0.6704 ± 0.02711 | 0.6104 ± 0.02931 | 0.6392 ± 0.02566 | 0.6657 ± 0.03950 | 0.6350 ± 0.03738 | 0.6220 ± 0.05480 | 0.7101 ± 0.03889 |
